# Modanovo: A Unified Model for Post-Translational Modification-Aware de Novo Sequencing Using Experimental Spectra from In Vivo and Synthetic Peptides

**DOI:** 10.1101/2025.09.12.675784

**Authors:** Daniela Klaproth-Andrade, Yanik Bruns, Wassim Gabriel, Christian Nix, Valter Bergant, Andreas Pichlmair, Mathias Wilhelm, Julien Gagneur

## Abstract

Post-translational modifications (PTMs) play a central role in cellular regulation and are implicated in numerous diseases. Database searching remains the standard for identifying modified peptides from tandem mass spectra, but is hindered by the combinatorial expansion of modification types and sites. De novo peptide sequencing offers an attractive alternative, yet existing methods remain limited to unmodified peptides or a narrow set of PTMs. Here, we curated a large dataset of spectra from endogenous and synthetic peptides from ProteomeTools spanning 19 biologically relevant amino acid-PTM combinations, covering phosphorylation, acetylation, and ubiquitination. We used this dataset to develop Modanovo, an extension of the Casanovo transformer architecture for de novo peptide sequencing. Modanovo achieved robust performance across these amino acid-PTM combinations (median area under the precision-coverage curve 0.92), while maintaining performance on unmodified peptides (0.93), nearly identical to Casanovo (0.94). The model outperformed π-PrimeNovo-PTM and showed increased precision and complementarity to the database search tool MSFragger. Robustness was confirmed across independent datasets, particularly at peptide lengths frequently represented in the curated dataset. Applied to a phosphoproteomics dataset from monkeypox virus-infected cells, Modanovo recovered numerous confident peptides not reported by database search, including new viral phosphosites supported by spectral evidence, thereby demonstrating its complementarity to database-driven identification approaches. These results establish Modanovo as a broadly applicable model for comprehensive de novo sequencing of both modified and unmodified peptides.

## Introduction

Post-translational modifications (PTMs) enhance the functional diversity of proteins by altering amino acid side chains after translation (1). These chemical modifications are relevant to protein activity, localization, and interactions, and are often implicated in disease (2). As a result, identifying PTMs is a central goal in proteomics and is essential for understanding many biological processes and diseases (3).

Bottom-up proteomics, typically using liquid chromatography-tandem mass spectrometry, is the primary method for large-scale protein and PTM identification (4,5). Here, proteins are enzymatically digested into peptides, which are first analyzed in a first mass spectrometry scan to measure their mass-to-charge (m/z) ratios (6–8). Selected precursor ions are then isolated and fragmented, and the resulting fragment ions are measured in a second scan, producing a tandem mass spectrum (9,10). Peptide sequences can, in principle, be inferred from mass differences between fragment ions (11,12), but this remains challenging due to spectral noise, missing peaks, co-fragmented peptides, and ambiguity in ion series assignment. Consequently, peptide identification, which serves as the foundation for subsequent protein inference and quantification, is typically performed through database searching (13,14). Here, experimental spectra are matched against theoretical spectra generated from the reference proteome of the organism of interest (15,16). However, including PTMs in the context of database searching exponentially increases the search space, making exhaustive searches increasingly difficult (17).

An alternative or complementary avenue to database search is de novo peptide sequencing (DNPS), which bypasses the reference database entirely, enabling the discovery of novel peptides, rare PTMs, and proteins from unsequenced organisms (18). Recent DNPS tools, including transformer-based models like Casanovo (19–21), have achieved strong results on benchmark datasets (22). However, identifying post-translationally modified peptides remains a major challenge. Casanovo and most of its peer-reviewed successor models (23–29) support only the seven amino acid-PTM combinations defined in the first release of the MassIVE-KB human spectral libraries (30), which include N-terminal modifications (acetylation, carbamylation, loss of ammonia), deamidation, and oxidation. In contrast, π-PrimeNovo (25) expands PTM coverage by fine-tuning on additional PTMs one at a time. It was trained separately for each of the PTM types contained in a dataset covering multiple PTMs (31), demonstrating flexibility through multiple single-PTM models, though the trained weights for all models have not been released publicly. A phosphorylation-specific π-PrimeNovo model trained on the 2020-Cell-LUAD dataset (32) has been released publicly. Although this single-PTM approach increases PTM diversity, it requires training individual models for each PTM, which is limited by the scarcity of large, diverse PTM datasets and the substantial computational costs of per-PTM training. Furthermore, in cases where multiple PTM types are of interest, each corresponding model must be run separately, and the user must determine which identification to trust, resulting in an approach that is user-biased and less practical for broad or ambiguous modification searches.

To the best of our knowledge, DNPS models that can simultaneously identify a broad spectrum of PTM types within a single unified framework while maintaining robust performance on unmodified peptides are lacking. This limitation partly arises from the scarcity of large-scale, high-quality experimental datasets covering diverse PTMs. Here, we compiled a sequence-annotated dataset of tandem mass spectra including unmodified peptides and 19 amino acid-PTM combinations, drawing from the MassIVE-KB spectral libraries (30) and the MULTI-PTM dataset, which is part of PROSPECT-PTM (33) and ProteomeTools (34). Using this resource, we developed Modanovo, a unified DNPS model built on the Casanovo architecture and expanding its PTM coverage by 12 amino acid-PTM combinations. Modanovo supports 39 tokens for canonical and modified residues, enabling broad PTM coverage without training separate models and while maintaining strong performance on unmodified peptides. Importantly, Modanovo identifies ubiquitinated, acetylated, and phosphorylated peptides, among others, reflecting PTMs with broad roles in protein regulation and cellular function. We demonstrate the utility of Modanovo by analysing human foreskin fibroblast (HFF) cells infected with monkeypox virus (MPXV), revealing relevant new phosphorylated peptides missed by conventional database search.

## Experimental Procedures

### Experimental Design and Statistical Rationale

The primary objective of this study was to develop and evaluate Modanovo, a de novo peptide sequencing (DNPS) model capable of handling datasets containing 19 amino acid-post-translational modification (PTM) combinations. A detailed description of the data processing workflow and model architecture is provided in subsequent sections. Modanovo was trained on experimental spectra from in vivo experiments from the MassIVE-KB (v1) human spectral libraries (30) and synthetic peptides from the MULTI-PTM dataset (33). Its generalization ability was evaluated on three independent publicly available datasets: 21 PTMs (31), MassIVE-KB (v2), and a dataset of MPXV-infected cells (35). To ensure strict separation of peptide sequences, peptide-spectrum matches (PSMs) were partitioned into training, validation, and test sets following the non-overlapping peptide splits established in the Casanovo study (19). Model performance was assessed using standard metrics for DNPS, including peptide-level precision-coverage curves and their corresponding area under the curve on held-out data. Benchmarking was performed against Casanovo, π-PrimeNovo (36), and MSFragger (37). No biological replicates were generated, as the study leveraged existing large-scale repositories.

For reproducibility, the source code is publicly available at: https://github.com/gagneurlab/Modanovo. Trained model weights and data splits are available at 10.57967/hf/6452 (38) and 10.57967/hf/6451 (39).

### Datasets

#### MassIVE-KB (v1) dataset

For model development, we downloaded a subset of the MassIVE knowledge base (MassIVE-KB) peptide spectral libraries (30) consisting of ∼30 million PSMs, which were also used for training Casanovo (19,20). These “high-quality” PSMs were identified by applying a very strict PSM-level false discovery rate (FDR) filter and selecting at most 100 PSMs for each precursor charge and modified peptide combination (30). The dataset includes both unmodified peptides and peptides bearing variable PTMs at specific residues. The variable PTM-residue combinations were: methionine oxidation (Unimod ID: 35), deamidation of asparagine and glutamine (Unimod ID: 7), N-terminal acetylation (Unimod ID: 1), N-terminal carbamylation (Unimod ID: 5), and N-terminal loss of ammonia (Unimod ID: 385). In addition, all cysteine residues were treated as fixed modified with carbamidomethylation.

We considered the same peptide-disjoint training, validation, and test sets that were used to develop Casanovo. From these training, validation, and test sets, we randomly selected a subset (∼9%) of ∼2.1M, ∼0.2M, and ∼0.2M PSMs, respectively, to be used for training and evaluation purposes of our model. Retaining a representative proportion of unmodified peptides and those PTMs unique to this dataset was intended to prevent forgetting during fine-tuning, while subsampling reduced computational cost and emphasized the additional PTMs not included in this dataset.

#### MULTI-PTM dataset

In the absence of large in-vivo datasets covering a wide range of PTMs for model development, we complemented our training data with a subset from PROSPECT-PTM (33), part of the ProteomeTools project (Zolg et al. 2017). This “MULTI-PTM dataset” comprises high-quality tandem mass spectra of synthetic peptides with 14 distinct variable PTM-amino acid combinations and was used for model training and evaluation. All modifications in this dataset are precisely localized, and each peptide contains at most one variable PTM in addition to methionine oxidation and the fixed carbamidomethylation of cysteine. The included PTM-residue combinations are: methionine oxidation; N-terminal and lysine acetylation (Unimod ID: 1); arginine citrullination (Unimod ID: 7); monomethylation of lysine and arginine (Unimod ID: 34); phosphorylation of serine, threonine, and tyrosine (Unimod ID: 21); lysine ubiquitinylation (Unimod ID: 121); pyroglutamate formation (pyro-Glu) from glutamic acid and glutamine (Unimod IDs: 27, 28); and O-GalNAc and O-GlcNAc modifications of serine and threonine (Unimod ID: 43).

We restricted the dataset to PSMs from higher-energy collisional dissociation (HCD; beam-type collision-induced dissociation) spectra and to those analyzed with an Orbitrap mass analyzer. Moreover, we randomly subsetted the dataset to contain at most 100 instances of the same unmodified peptide and at most 200 instances of the same modified peptide. This resulted in a total of over 8 million PSMs, which were randomly divided into training, validation, and test sets comprising approximately 90%, 5%, and 5% of the data, respectively. The defined sets are disjoint at the peptide level and follow the same split strategy as used in Casanovo. Moreover, the modified and unmodified counterparts of each peptide sequence were always placed in the same split.

#### 21-PTM dataset

We obtained the raw files and MaxQuant (40) identification results for the 21-PTM dataset (31), part of the ProteomeTools project (34), which contains synthetic peptides with 21 distinct amino acid-PTM combinations, from the PRIDE repository PXD009449. We restricted the dataset to PSMs derived from HCD fragmentation acquired on Orbitrap mass analyzers. We subset the dataset to contain only PTMs present in the MULTI-PTM and MassIVE-KB (v1) datasets, resulting in a total of six amino acid-PTM combinations. Nevertheless, we continue to refer to it as the “21-PTM dataset,” consistent with its common usage in the field.

#### MassIVE-kb (v2) dataset

We downloaded the second release of the MassIVE Knowledge Base (MassIVE-KB v.2024-05-24, (30)), which extends the variable PTMs cataloged in the first release by including phosphorylation and ubiquitination. Consistent with the download process for the initial release, we applied stringent quality controls by selecting “high-quality” PSMs and limited the dataset to a maximum of 100 PSMs per unique combination of precursor charge and peptide sequence. For this study, we further restricted the dataset to PSMs corresponding exclusively to post-translationally modified peptides, resulting in ∼150 thousand PSMs.

#### Monkeypox virus (MPXV) dataset

We obtained the raw files and MaxQuant identification files for the full proteome and phosphorylation-enriched datasets of human foreskin fibroblasts (HFF) cells infected with monkeypox virus (MPXV) from the PRIDE repositories PXD040811 and PXD040889 (35). We discarded decoy and secondary peptides so that each experimental spectrum is attributed to at most one peptide sequence, namely the one with the highest score.

### Identification of post-translationally modified peptides with Modanovo

#### Fine-tuning of a transformer model for peptide prediction from tandem mass spectra

We based Modanovo on the Casanovo transformer architecture, which employs a sequence-to-sequence framework to predict peptide sequences directly from tandem mass spectra. The model formulates peptide sequencing as a next-token prediction task, where each token represents either a canonical amino acid or a PTM-amino acid combination.

The original Casanovo vocabulary consisted of 28 tokens, including the special stop token and 7 amino acid-PTM combinations. To accommodate an expanded vocabulary of 40 tokens reflecting the inclusion of 12 new PTM-amino acid combinations as distinct tokens, we increased the dimensions of the input embedding layer and the output projection layer accordingly. The model with extended token vocabulary was initialized with pre-trained weights from Casanovo (v4.0.0), originally trained on the MassIVE-KB (v1) dataset comprising primarily unmodified peptides. Embeddings for the new tokens were initialized by averaging those of canonical amino acids, providing a meaningful initialization that facilitates efficient learning. We fine-tuned the entire model end-to-end on the combined training dataset containing spectra from both unmodified and modified peptides, enabling the model to learn fragmentation patterns associated with a broader spectrum of PTMs.

Training was conducted with a learning rate of 1 · 10^−5^, a dropout probability of 0.1, a batch size of 32, and a maximum of 12 epochs. Model performance was monitored on the validation set to mitigate overfitting, and the model exhibiting the lowest validation loss was selected as the final version for all subsequent analyses. All other training strategies and preprocessing parameters were kept consistent with the default settings established in Casanovo.

### Evaluation and alternative methods

#### Peptide prediction score

For ranking and evaluation, each PSM was associated with a confidence score. For Modanovo and Casanovo, a peptide-level score was calculated as the mean of the amino acid scores, taken directly from the model’s softmax output at each decoding step. To stabilize residue-level confidences, each amino acid score was averaged with the overall peptide score. If the predicted sequence did not match the precursor mass within the specified tolerance, a penalty of -1 was applied to the peptide-level score.

For π-PrimeNovo, peptide confidence was defined as the mean predicted probability across all residues in a sequence, as reported by the non-autoregressive transformer. For MSFragger, peptide scores were taken from the search engine’s reported spectral matching score (Hyperscore).

#### Evaluation metrics

During model training and evaluation, isoleucine and leucine were treated as indistinguishable due to their identical masses. Additionally, pyro-Glu formation from glutamic acid and glutamine is treated as a single PTM-amino acid combination, reflecting the chemical equivalence of the resulting pyroglutamate residue.

We evaluated the model performance of each DNPS tool using peptide-level precision-coverage curves. For each spectrum, we compared the peptide predicted by a tool to the corresponding peptide identified by a database search engine at a specified FDR threshold. The score refers to the confidence score assigned by the tool. A PSM was considered correct if it exactly matched the database-identified peptide, allowing for substitutions mentioned above. PSMs with a ground truth database identification but no prediction were considered incorrect and assigned the lowest possible score. Precision and coverage were computed across varying score thresholds *t* among all PSMs identified by the database search engine, and were defined as:

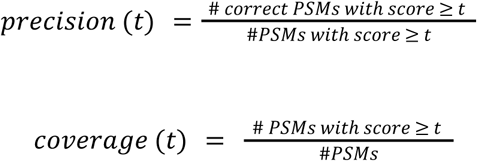

The area under the precision-coverage curve (AUPCC) was then computed using the trapezoidal method as implemented in the auc function of the scikit-learn package (41). In addition to the AUPCC, we often reported the final precision (at coverage 1) for each tool, which is equivalent to the peptide recall obtained by the tool in comparison to the ground truth identifications. To compute peptide-level precision-coverage curves for a given PTM-amino acid combination, we include all ground truth peptides that contain that specific modification, regardless of whether additional modifications are also present. For example, the ground truth peptide “PEPT[+79.966]IDEK[+14.016]” contributes to both the peptide-level precision-coverage curves for phosphorylated threonine (T[+79.966]) and monomethylated lysine (K[+14.016]).

#### Peptide alignment

Peptide alignments were obtained by running blastp (version 2.12.0+) against sequences of the reviewed human proteome, including isoforms (Uniprot, Taxon ID 9606) and Monkeypox virus proteins (GenBank: ON563414.3), downloaded from the PRIDE repository PXD040811. As a scoring matrix, we used the identity matrix, modified such that leucine and isoleucine were considered equivalent. All other blastp settings were set to their default values. We restricted the output of blastp to at most one hit per queried peptide sequence. If multiple hits were returned by the search, we selected the hit with the lowest e-value. We defined a query peptide to be a perfect alignment if the peptide is identical to the target peptide (except for differences between leucine and isoleucine).

#### Validation with Prosit

We obtained Prosit (42) spectrum predictions through Koina (43) for unmodified peptide identifications in the MPXV dataset using Prosit. Peptides longer than 30 residues, shorter than 7 residues, or with a charge state exceeding 6 were excluded. Prosit-predicted spectra for a given peptide sequence were matched to the corresponding experimental spectra by aligning each Prosit-predicted peak to the nearest experimental peak within a 20 ppm tolerance window, if present. The normalized spectral angle (SA) between the predicted and experimental spectra is defined as:

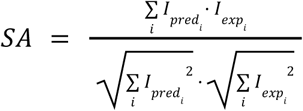

where 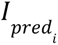 and 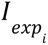 denote the intensities of the i-th matched peak in the predicted and experimental spectra, respectively. Both experimental and theoretical intensities were normalized using base-peak normalization before SA computation. If no experimental peak was found within the tolerance window to match an expected peak, the intensity 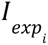 was set to zero. The SA ranges from 0 to 1, with values closer to 1 indicating greater similarity.

#### Empirical precision estimates on the MPXV dataset

We obtained empirical precision estimates on the phosphorylation-enriched and full proteome samples of the MPXV dataset by leveraging PSMs identified with MaxQuant as a reference. For each score threshold of Modanovo, we calculated the proportion of PSMs that could be matched to a MaxQuant peptide identification. This proportion was used as an empirical estimate of the precision at that threshold. By scanning across thresholds, we determined the score cutoffs that corresponded to target precision levels of 80%, 90%, and 95% for the full proteome and phospho-enriched samples separately. These empirically derived cutoffs were then applied to the set of PSMs without MaxQuant peptide identifications, enabling the extension of the empirical precision estimates to all identifications.

#### MPXV H5 structure modeling with AlphaFold and electrostatic surface potential analysis

In silico prediction of the structure of the MPXV H5 dimer was performed using the colab version of AlphaFold 2.3.1 (44) in the multichain mode using default parameters. The electrostatic surface potential of the modeled structure of the MPXV H5 dimer was calculated using the PyMOL plugin APBS electrostatics. Molecular graphics depictions were produced with the PyMOL software (45).

#### Alternative methods

##### Casanovo

We downloaded the Casanovo (v4.0.0) model weights from https://github.com/Noble-Lab/casanovo/releases/tag/v4.0.0. Casanovo was trained on the MassIVE-KB (v1) dataset, and we obtained the corresponding training, validation, and test splits in April 2024 from https://noble.gs.washington.edu/~melih/mskb_casanovo_splits.zip.

##### π-PrimeNovo-PTM

We cloned the π-PrimeNovoPTM (36) code from https://github.com/PHOENIXcenter/pi-PrimeNovo/tree/main/pi-PrimeNovo-PTM and downloaded the model weights fine-tuned for phosphorylation. This model was trained on the 2020-Cell-LUAD dataset, which focuses on human lung adenocarcinoma and includes 103 LUAD tumor samples along with their matched non-cancerous adjacent tissues (32). This π-PrimeNovo-PTM model represents phosphorylation using a dedicated token “B” with a corresponding mass of 79.9663. For evaluation, we accepted predicted sequences as correct provided the phosphorylation was localized to the correct serine, threonine, or tyrosine residue, irrespective of whether the “B” token preceded or followed the residue.

##### MSFragger

We ran MSFragger (v4.1 (37)) on the MULTI-PTM dataset through the FragPipe interface (v22.0) using a closed search configuration. Carbamidomethylation of cysteine was specified as a fixed modification, and all PTM-amino acid combinations covered by Modanovo were explicitly specified as variable modifications in the search parameters. Spectra were searched against a forward and reverse version of the reviewed human proteome, including isoforms (UniProt, Taxon ID 9606, containing 20,596 proteins and downloaded November 17, 2023). The search was performed using tryptic digestion with precursor and fragment mass tolerances set to 20 ppm. PSM-level FDR filtering was performed using Percolator with an FDR threshold of 0.01. Peptide-level and protein-level FDR thresholds were set to 1 to avoid additional filtering beyond the spectrum level.

## Results

### A dataset for developing de novo PTM identification models

We compiled a dataset combining a subset of the MassIVE-KB human spectral libraries (v1), consisting mostly of unmodified peptides and 7 amino acid-PTM combinations, with a curated subset of spectra from MULTI-PTM (part of PROSPECT-PTM (33) and the ProteomeTools project (34)). The latter includes synthetic peptides modified with a range of biologically relevant post-translational modifications (PTMs) such as phosphorylation, acetylation, ubiquitylation, and methylation, which were crucially missing in MassIVE-KB (v1). Due to the scarcity of large, well-annotated experimental datasets covering diverse PTMs, this combination allowed us to leverage both data from in vivo experiments as well as from synthetic peptides to improve model generalization. In total, this combined dataset, denoted as development dataset throughout this manuscript, consisted of approximately 11 million PSMs spanning 20 canonical amino acids and 19 distinct PTM-amino acid combinations, 12 of them not being covered by Casanovo and most of its successor models (Fig. 1A, B, Supplementary Table 1), enabling comprehensive learning and evaluation across a broad spectrum of peptide modifications.

**Figure 1.**
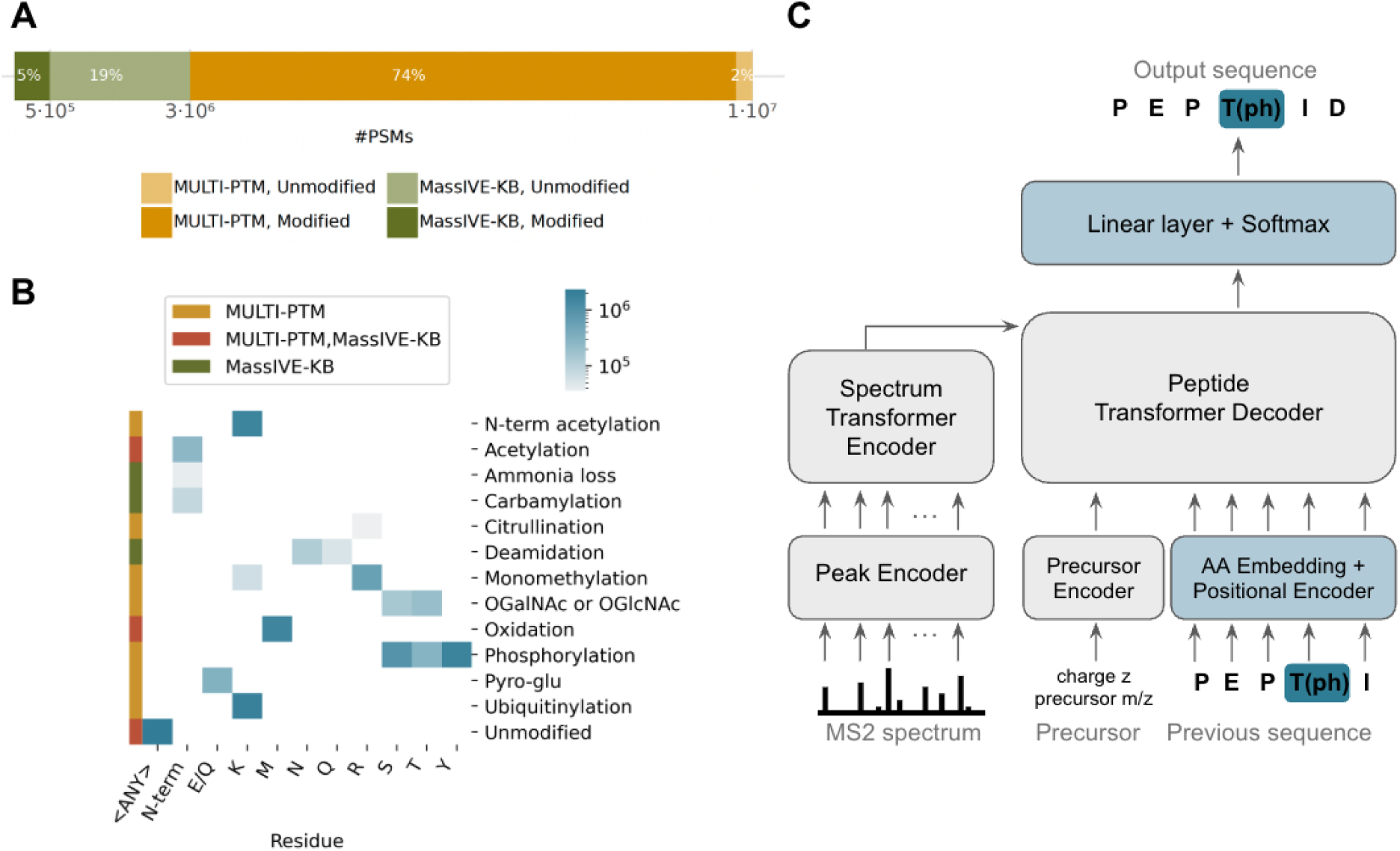
**Modanovo identifies post-translationally modified peptide sequences using a mixture of spectra from in vivo experiments and synthetic peptides for model development**. **A**, Number of peptide-spectrum matches (PSMs) for modified and unmodified peptides used for model development from two different data sources: the MassIVE-KB (v1) human spectral libraries (dark and light green) and from a subset of PROSPECT-PTM (MULTI-PTM, dark and light mustard). **B**, Heatmap showing the number of PSMs used for model development across PTM types and modified residues. Rows correspond to PTM types, columns to residues. The side color strip indicates the data source (MULTI-PTM, MassIVE-KB, or both). The heatmap is colored on a logarithmic scale, with darker blue shades representing higher PSM counts. **C**, An autoregressive transformer encoder and decoder architecture based on Casanovo allows the identification of post-translationally modified peptides directly from tandem mass (MS2) spectra. The model is trained starting with weight initialization from Casanovo’s pre-trained weights. The model components for the amino acid (AA) embeddings and final linear layer (in blue) are expanded to allow the modification of new post-translationally modified residues.

### Modanovo extends Casanovo to 12 new amino acid-PTM combinations

Based on this development dataset, we developed Modanovo, a transformer-based model designed to identify modified peptides directly from tandem mass spectra. To enable this, we fine-tuned Casanovo’s transformer architecture (Fig. 1C), initializing the model with weights from a Casanovo model previously trained on MassIVE-KB (v1). To accommodate the expanded vocabulary arising from PTM-amino acid combinations treated as distinct tokens, we adjusted the shapes of the input embedding layer and the final linear projection layer (Methods). The new parameters corresponding to the expanded token set were initialized by averaging the weights of existing tokens in their respective layers, providing a meaningful starting point rather than random initialization. We then fine-tuned the entire model end-to-end, allowing it to retain knowledge of canonical peptide fragmentation patterns while adapting to the variability introduced by PTMs.

### Modanovo confidently identifies modified synthetic peptides

We first assessed whether Modanovo’s performance remained comparable to that of Casanovo on unmodified peptides and the PTMs covered by Casanovo, which served as the starting point of our fine-tuning approach for PTM expansion. Specifically, we evaluated both models on the test set of the MassIVE-KB (v1) dataset, which was also used to originally develop and evaluate Casanovo and primarily consists of unmodified peptides. We found that Modanovo achieves nearly identical performance to Casanovo overall, with an area under the precision-coverage curve (AUPCC) of 0.93 versus 0.94 (on held-out data here and everywhere else, Supplementary Fig. S1A), indicating that it remains well suited for identifying unmodified peptides. In terms of AUPCC across specific PTM-amino acid combinations, Modanovo shows strong agreement with Casanovo (Supplementary Fig. S1B), with only minor variations observed across individual PTM categories. Slightly larger performance differences are seen for deamidation and N-terminal loss of ammonia. The reduced performance for peptides with N-terminal ammonia loss may reflect added complexity introduced by the inclusion of pyroglutamate and citrullination modifications, which result in similar mass shifts and may interfere with model discrimination in this mass range.

Having confirmed that Modanovo maintains strong performance on unmodified peptides from in vivo experiments and the set of PTMs covered by Casanovo, we evaluated Modanovo on the MULTI-PTM proportion of the development dataset. The MULTI-PTM dataset contained 12 distinct PTM-amino acid combinations, which did not overlap with those covered in Casanovo. Across these distinct PTM-amino acid combinations, Modanovo achieved a median AUPCC of 0.92 ([0.70, 0.96] 95% confidence interval) and a median final peptide precision of 0.68 ([0.48, 0.74] 95% confidence interval), measured against ground truth peptide sequences reported by PTM-specific MaxQuant (40) searches (Fig. 2A). Comparably, on unmodified peptides from the development dataset, Modanovo attained a median AUPCC of 0.93 and a final peptide precision of 0.71 (Fig. 2A). Particularly, phosphorylated peptides are accurately identified by Modanovo, with AUPCC values of 0.96, 0.93, and 0.94, and final precision values of 0.78, 0.71, and 0.72 for sequences containing phosphorylated serine (S[+79.966]), threonine (T[+79.966]), and tyrosine (Y[+79.966]) residues, respectively. Comparable performance is observed for peptides bearing acetylation (K[+42.011] and [+42.011]-), ubiquitination (K[+114.043]), and citrullination (R[+0.984]). Notably, Modanovo achieves an AUPCC of 0.96 for sequences containing pyroglutamate residues (E[-18.011] and Q[-17.027]). This performance, which surpasses that observed for unmodified peptides, may be due to the model successfully learning that pyroglutamate formation is restricted to the first residue position in a peptide sequence.

**Figure 2.**
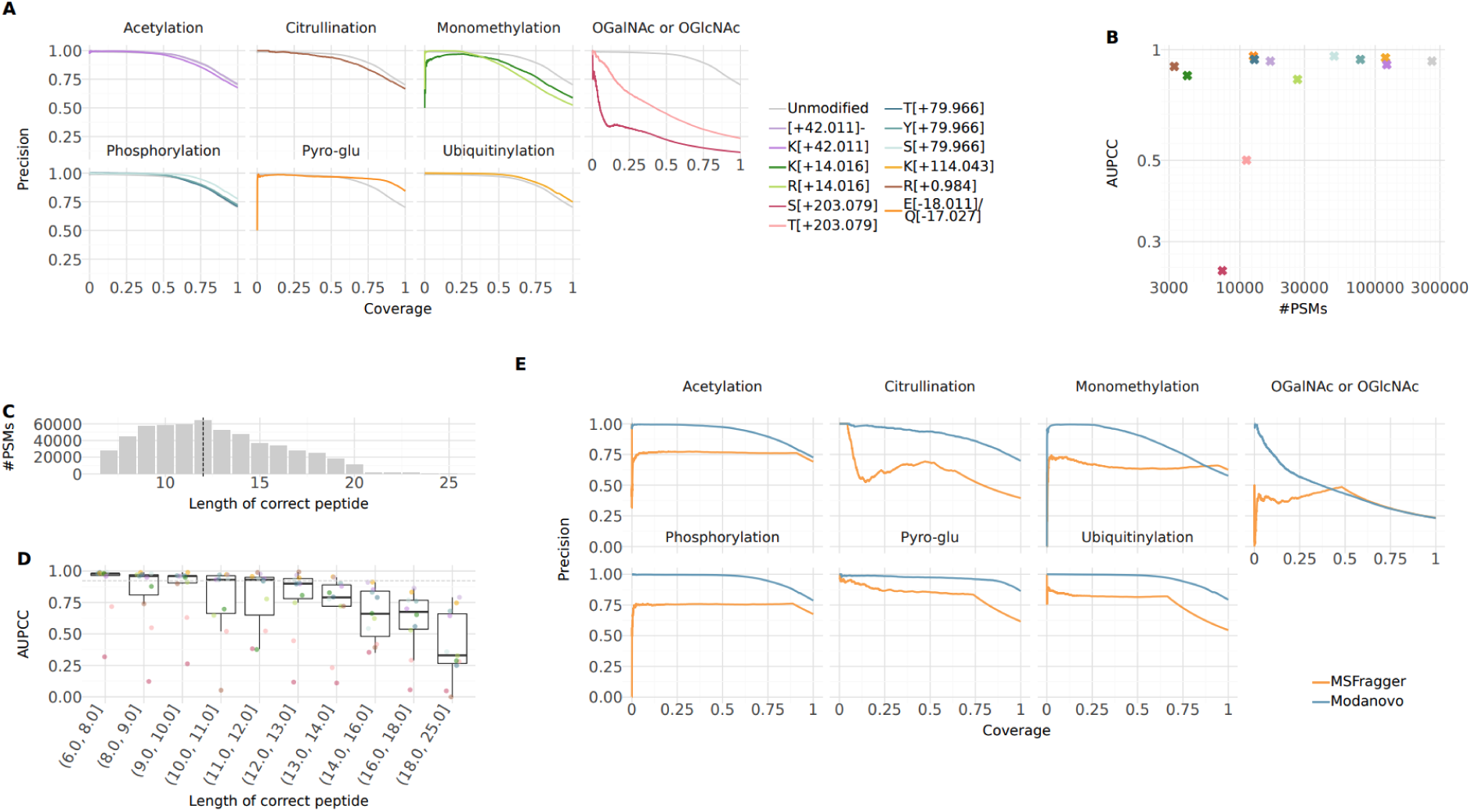
**Model performance on the test set of the development dataset**. **A**, Precision-coverage curves at the peptide level for the 12 new PTM-amino acid combinations covered by Modanovo, as well as methionine oxidation. PTM types (Acetylation, Citrullination, Monomethylation, OGalNAc/OGlcNAc, Oxidation, Phosphorylation, Pyro-glu, and Ubiquitination) are shown in the different panels, and colors represent different PTM-residue combinations. Performance for unmodified peptides (light grey) is shown in each panel for comparison. Precision-coverage curves for the remaining PTM-amino acid combinations, which were covered by Casanovo before, are found in Supplementary Fig. S1 **B**, Area under the precision-coverage curve (AUPCC) against the number of peptide-spectrum matches (PSMs) per PTM-residue combination on the test set, using the same color scheme as in panel A. Statistical significance was assessed using Spearman correlation (ρ = 0.45, p = 0.13). **C,** Distribution of peptide sequence lengths in the test set of the MULTI-PTM dataset. The dashed vertical line marks the median peptide length. **D,** AUPCC at the peptide level across peptide length bins, evaluated on the test set. Each point represents a PTM-amino acid combination, using the same color scheme as in panel A. Bins were constructed to contain approximately equal numbers of PSMs. The dashed grey line indicates the AUPCC across all peptide lengths. **E,** Precision-coverage curves at the peptide level comparing Modanovo (blue) to MSFragger (claimed 1% FDR, orange) on the test set of the MULTI-PTM dataset, faceted by the different PTM types. MSFragger often does not propose a peptide for a given spectrum. These are ranked last and cause the hyperbole sections on the higher coverage range.

In contrast to the consistently high performance observed across most PTM types, Modanovo shows reduced performance on peptides containing O-GalNAc and O-GlcNAc modifications of serine and threonine (AUPCC of 0.25 and 0.5). Perhaps, this is partly because these glycosylation events produce complex and often heterogeneous fragmentation patterns, with reduced fragment ion coverage and intensity that hinder reliable sequence reconstruction (46). Moreover, both modifications are encoded using the same token in the model due to their mass equivalence and equal Unimod identifier (47), despite their structural differences. The challenge is compounded by the limited number of examples for these glycopeptides in the dataset (Fig. 2B, ∼19,000 glycopeptides vs. a mean number of test PSMs per PTM type in the MULTI-PTM dataset of 66,109). Interestingly, when allowing modified peptide sequences to be considered correct despite shifts in the O-GalNAc/O-GlcNAc modification site between serine and threonine (e.g., treating PEPTI**S[+203.079]**ER and PEP**T[+203.079]**ISER equivalently), or between two serine or threonine residues, the AUPCC improved (from 0.25 to 0.58 for serine residues and from 0.50 to 0.70 for threonine residues, Supplementary Fig. S2), although it still fell short of the levels achieved for other PTM types. This improvement shows that while the model sometimes struggles to localize the modification to the exact residue, it more often correctly calls the modification at the peptide level and recovers the underlying unmodified peptide sequence. These shifts likely reflect the inherent difficulty of pinpointing the modification site when spectra lack clear site-determining fragment ions. Similarly, the somewhat decreased performance of monomethylated peptides can be partly attributed to disagreements in the localization sites between the ground truth and predicted peptide sequences. Allowing for different monomethylation sites increased the AUPCC from 0.83 to 0.85 for arginine residues and from 0.85 to 0.91 for lysine residues (Supplementary Fig. S2).

Overall, a moderate positive correlation was observed between the AUPCC and the number of PSMs in the test set, but the relationship was not statistically significant (Fig. 2B, Spearman’s ρ = 0.45, *p* = 0.13). This suggests that factors beyond dataset size, such as fragmentation behavior or the structural properties of specific PTMs, play an important role in model performance. Notably, some PTMs with relatively few examples, such as citrullination, still achieved competitive AUPCC values, while others with larger sample sizes, such as arginine monomethylation, perform more modestly. These findings indicate that PTM-specific learnability may outweigh the absolute amount of training data in determining predictive accuracy.

We reasoned that achieving high performance on longer peptides is more challenging, since every amino acid and its modifications must be correctly identified, the spectra of long peptides are often of lower quality, and errors accumulate due to the autoregressive nature of the transformer model. Therefore, we evaluated the impact of peptide length on Modanovo performance. Peptides on the MULTI-PTM subset were 12 residues long in median (Fig. 2C). Modanovo achieved improved or comparable median AUPCC across PTM-residue combinations, relative to the overall median AUPCC (0.92), for peptides up to 13 residues in length (Fig. 2D). However, model performance declined for longer peptides. For example, the AUPCC for sequences containing phosphorylated serine dropped from 0.93 to 0.78 when comparing peptides of length 12-13 to those of length 13-14. Similar observations were made on the Massive-KB (v1) dataset for both Modanovo and Casanovo, whose performance decreased for peptides longer than the median length of 14 (Supplementary Fig. S3). These findings suggest that Modanovo performs most reliably within the peptide length range it was most frequently exposed during training.

Among peer-reviewed state-of-the-art DNPS tools claiming to expand PTM coverage beyond Casanovo’s vocabulary, publicly available model weights are limited to versions supporting phosphorylation only. Therefore, we compared Modanovo to the publicly available π-PrimeNovo-PTM (25) model, which extends Casanovo’s token vocabulary to enable direct prediction of phosphorylated residues, on phosphorylated peptides from the development dataset. Modanovo substantially outperformed π-PrimeNovo-PTM on all phosphorylated sequences of the dataset (Supplementary Fig. S4).

Having established Modanovo as a DNPS model covering 12 new amino acid-PTM combinations, we next asked whether it could serve as a useful complementary tool to state-of-the-art database search approaches. For this, we first compared the performance of Modanovo on the 12 new amino acid-PTM combinations to that of MSFragger (37), applying a 1% PSM-level FDR. As a realistic database search reflecting the application setting, in which multiple PTMs are of interest and the ground truth is unknown, we ran MSFragger against the full human proteome, allowing for the same set of possible modifications as Modanovo (Methods). We note that this database search setup differs substantially from the one initially used to establish high-quality ground truth annotations (33), where targeted searches were performed with MaxQuant, each restricted to the relevant modification and using a database limited to the synthesized peptides. Overall, Modanovo outperformed MSFragger in identifying modified peptides across all PTM types (Fig. 2E), highlighting its strong ability to identify correct PSMs even at high precision levels.

### Modanovo generalizes to independent datasets

To evaluate Modanovo’s generalizability, we applied it to two distinct datasets: the “21-PTM dataset” (31), consisting of modified synthetic peptides, and a modified-only subset of the latest release of MassIVE-KB (v2), which, in contrast to the first release, included phosphorylated, acetylated, and ubiquitinated residues. We focused our assessment on the 12 new amino acid-PTM combinations covered by Modanovo, resulting in 6 and 5 amino acid-PTM combinations from the 21-PTM and MassIVE-KB (v2) datasets. For consistency with common usage in the field, we still refer to the first dataset as the “21-PTM dataset”, even though only a subset of PTMs was considered here.

While the performance generally decreased compared to the development dataset, they remained strong across a diverse range of amino acid-PTM combinations (Fig. 3A, Supplementary Fig. S5), though the quality of identification varied depending on the dataset and modification. For instance, the model identified peptides with lysine ubiquitylation (K[+114.043]) with an AUPCC of 0.93 and 0.82 on the 21-PTM and MassIVE-KB (v2) datasets, respectively, compared to 0.95 on the development dataset. Similarly, it reached an AUPCC of 0.89 for lysine acetylation (K[+42.011]) on the 21-PTM dataset, compared to 0.91 for the development dataset. The AUPCC was generally lower on MassIVE-KB (v2) for the remaining PTM types (acetylation and phosphorylation, Fig. 3A), with the most pronounced drop observed for phosphorylation (mean AUPCC: 0.94 vs. 0.65). However, this decrease in performance was largely explained by differences in peptide length distributions: MassIVE-KB (v2) contains a greater proportion of longer peptides than the 21-PTM and development datasets (median lengths: 19, 14, and 12, Supplementary Fig. S6). When controlling for peptide length, the AUPCC remained comparable between the three datasets (Fig. 3B), indicating that Modanovo’s learned representations generalize well to in vivo-derived spectra. This consistent performance at matched lengths supports the conclusion that Modanovo is capable of accurately identifying modified peptides from complex biological samples, despite being trained primarily on data from synthetic peptides.

**Figure 3.**
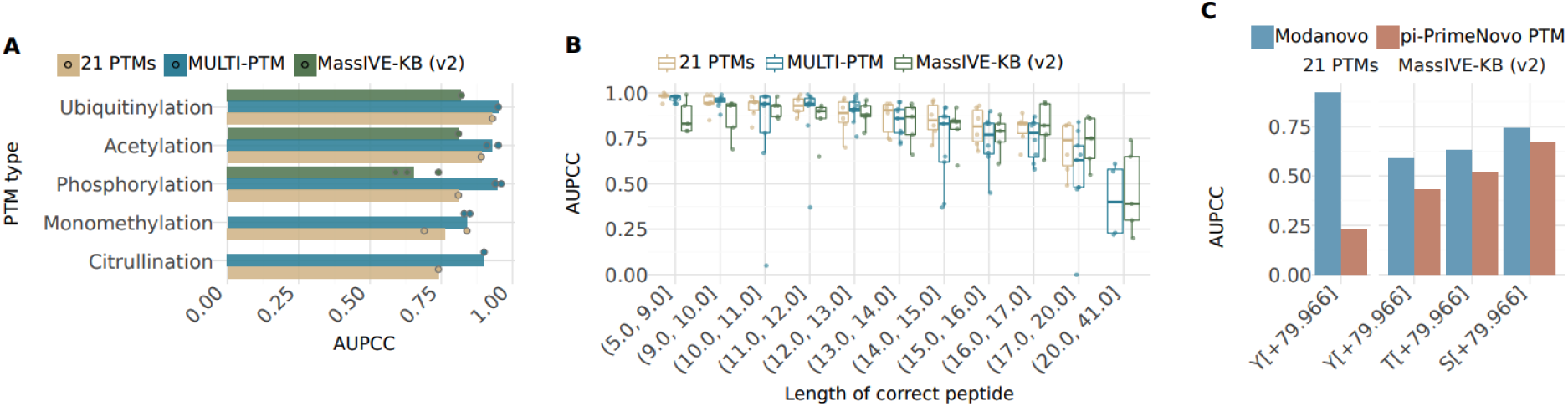
**Model performance on alternative datasets and comparison to other methods**. **A**, Area under the precision-coverage curve (AUPCC) for the different PTM types contained in the three different datasets, MassIVE-KB (v2, green), MULTI-PTM (blue), and 21 PTMs (light brown). Data points represent individual PTM-residue combinations, restricted to the overlap between amino acid-PTM combinations present in the MassIVE-KB or 21-PTM dataset and new 12 amino acid-PTM combinations introduced in Modanovo. **B,** AUPCC across peptide length bins for the three different datasets, MassIVE-KB (v2, green), MULTI-PTM (blue), and 21 PTMs (light brown). Each point represents an amino acid-PTM combination. Bins were constructed to contain approximately equal numbers of PSMs. **C**, AUPCC for the different phosphorylated residues by Modanovo (blue) compared to π-PrimeNovo-PTM (brown) on the MassIVE-KB (v2) and 21-PTM datasets.

Unlike the peptides in the development dataset, the 21-PTM dataset contains sequences with more than one PTM type (beyond methionine oxidation). To evaluate Modanovo’s ability to generalize to these more complex cases, which were not seen during training, we stratified performance by PTM-type combinations. The model was able to sequence multiply modified peptides for most combinations (Supplementary Fig. S7). For example, Modanovo achieved an AUPCC of 0.74 for peptides containing both acetylated and monomethylated residues (K[+42.011], R[+14.016]), which, while lower than the values observed for singly modified peptides (0.76 for monomethylation and 0.95 for acetylation), still indicates reasonable generalization to peptides carrying multiple co-occurring modifications despite being trained only on singly modified peptides (apart from optional methionine oxidation).

Phosphorylation site (P-site) localization remains a longstanding challenge in mass spectrometry-based proteomics. To assess Modanovo’s robustness to this ambiguity, we evaluated its ability to identify phosphorylated peptides in the MassIVE-KB (v2) and 21-PTM datasets while allowing for alternative P-site placements relative to those assigned by database searching. While performance remains unchanged on the 21-PTM dataset when accepting predictions with different P-sites, either on the same residue type or on a different phosphorylatable residue, it increased for the MassIVE-KB (v2) dataset (Supplementary Fig. S8). For example, among peptides with a phosphorylated tyrosine as the ground truth, the AUPCC improved from 0.74 to 0.79 when predictions placing the phosphate on serine or threonine residues were also considered correct. These results underscore Modanovo’s flexibility in recovering plausible phosphopeptides despite the inherent uncertainty of P-site assignment.

We further compared Modanovo to the publicly available π-PrimeNovo-PTM (25) model, which allows the prediction of phosphorylated residues. Consistent with the observations in the development dataset (Supplementary Fig. S4), Modanovo consistently achieved a higher AUPCC than π-PrimeNovo-PTM across all three phosphorylated residue types (Fig. 3C, Supplementary Fig. S9), with gains of 0.59 vs. 0.43 for tyrosine, 0.63 vs. 0.52 for threonine, and 0.74 vs. 0.67 for serine, on the MassIVE-KB (v2) dataset; and substantially outperformed π-PrimeNovo-PTM on the 21-PTM dataset. These results suggest that explicitly modeling PTM-amino acid combinations (e.g., S[+79.966]) as single tokens, rather than treating the PTM mass shift (e.g., [+79.966]) as an independent token that can appear anywhere in the sequence, may contribute to improved predictive performance. Combined with training on a larger and more diverse dataset containing several PTM types, this design choice could help better capture biologically plausible fragmentation patterns across different PTM types.

### Modanovo identifies phosphopeptides from Monkeypox virus-infected samples

Having established Modanovo as a DNPS tool capable of confidently identifying post-translationally modified peptide sequences from tandem mass spectra, we applied it to a time-resolved dataset of human foreskin fibroblast (HFF) cells infected with monkeypox virus (MPXV), comprising both full proteome and phosphoproteome measurements (35). This recently published dataset represents a compelling use case, as it captures the complex interplay of PTMs in both the virus and the host during the course of infection. Furthermore, DNPS approaches are particularly attractive in viral proteomics, since viral genomes frequently undergo mutations that can complicate database-driven peptide identification. The dataset was originally analyzed with the database search engine MaxQuant. Here, we evaluated the extent to which Modanovo provides complementary insights to this initial analysis.

MaxQuant identified a peptide sequence at 1% FDR for only 15% and 30% of the spectra in the phospho-enriched samples and full proteome samples, respectively. Considering these identifications as ground truth, Modanovo demonstrated strong performance on unmodified peptides and peptides containing only common modifications (N-terminal acetylation or methionine oxidation), achieving an AUPCC of 0.95 and 0.88 on phosphorylation-enriched and full proteome samples, respectively (Fig. 4A, Supplementary Fig. S10). This was consistent with the performance on the development dataset (AUPCC of 0.93) and further confirmed the robustness of the model. For peptides containing a single phosphorylated residue, the AUPCC decreased to 0.78. While this represents a drop in performance, it remained comparable to Modanovo’s performance on unmodified peptides and phosphopeptides of similar lengths in the development dataset (Fig. 2D, Supplementary Fig. S11). Hence, the lower performance on phosphopeptides in the MPXV dataset is attributable to a shift in peptide length distribution: the median sequence lengths were 16 and 17 amino acids for singly and multiply phosphorylated peptides, respectively (Supplementary Fig. S12), longer than those typically observed during training. As expected, model performance declined with increasing proportions of missing y-ion fragments (Supplementary Fig. S13). Notably, the model performance for singly phosphorylated peptides closely matched that of unmodified peptides when controlling for the proportion of missing y-ions (Supplementary Fig. S13). Overall, Modanovo maintained competitive performance when using MaxQuant identifications as ground truth, highlighting its ability to generalize to more complex datasets with longer peptides and variable fragment coverage.

**Figure 4.**
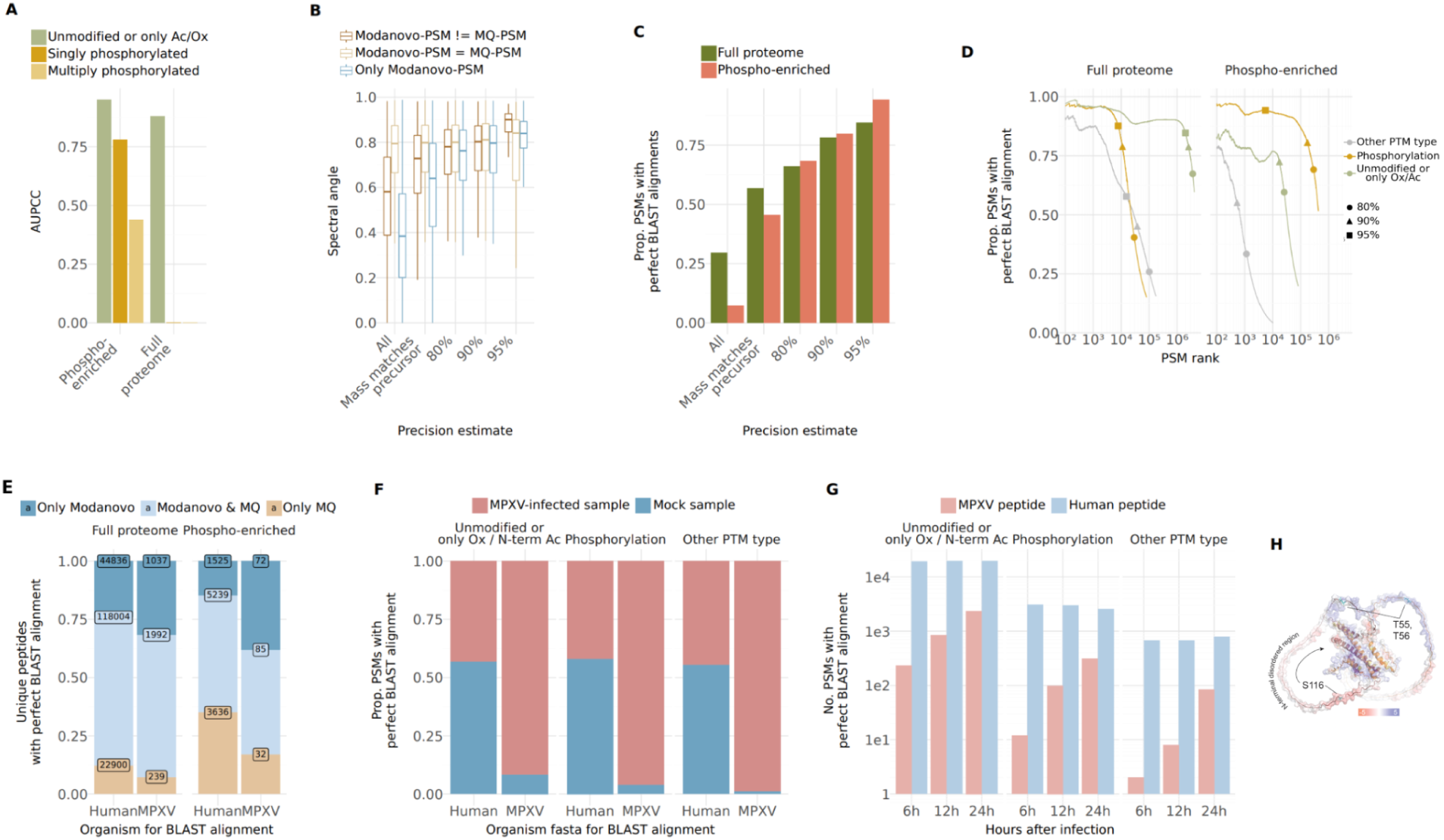
**Modanovo applied to a dataset of monkeypox virus (MPXV)-infected human foreskin fibroblast (HFF) cells**. **A,** Area under the precision-coverage curve (AUPCC) obtained with Modanovo’s predictions with MaxQuant peptides as ground truth sequences for comparison for the full proteome samples and phosphorylation-enriched samples with the MaxQuant sequence having multiple phosphorylated residues (light mustard), one phosphorylation residue (mustard), and no phosphorylation residues (green). **B,** Spectral angles obtained from the comparison of experimental spectra and Prosit-predicted spectra for Modanovo’s prediction for the whole set of predictions (“All”), for predictions with mass matching the precursor mass-to-charge and at different precision estimates (80%, 90% and 95%), and for Modanovo’s predictions matching the identified MaxQuant sequence (light brown), not matching the MaxQuant sequence (dark brown), and for spectra without a MaxQuant identification (blue). **C,** Proportion of peptide-spectrum matches (PSMs) with perfect BLAST alignments for the phosphorylation-enriched (terracotta) and full proteome (green) samples for all PSMs predicted by Modanovo (“All”), the PSMs predicted by Modanovo with calculated mass matching the precursor m/z (“Mass matches precursor”), and at different precision estimates (80%, 90%, 95%). **D,** Proportion of PSMs with perfect BLAST alignments and mass consistent with the precursor m/z for the phosphorylation-enriched (right) and full proteome (left) samples, sorted by Modanovo’s score for phosphorylated peptides (mustard), unmodified peptides, or peptides only containing methionine oxidation or N-terminal acetylation (green) and peptides containing other PTM types covered by Modanovo (grey). For clarity, the first 100 PSMs are omitted. **E,** Proportion of unique peptides with perfect BLAST alignment and mass consistent with the precursor m/z for the phosphorylation-enriched (right) and full proteome (left) samples obtained by querying against the human and MPXV proteome for sequences only identified by Modanovo (blue), Modanovo and MaxQuant (MQ, light blue), and only MaxQuant (brown). **F,** Proportion of PSMs with perfect BLAST alignments and mass consistent with the precursor m/z for the phosphorylation-enriched (right) and full proteome (left) samples obtained by querying against the human and MPXV proteome for MPXV-infected samples (terracotta) and mock samples (blue). **G,** Number of PSMs with perfect BLAST alignments and mass consistent with the precursor m/z for the phosphorylation-enriched (right) and full proteome (left) samples for samples measured at different hours post-infection for MPXV peptides (light terracotta) and human peptides (light blue). **H,** In silico predicted structure of the MPXV H5 dimer by AlphaFold (44) overlaid with electrostatic surface potential analysis of non-phosphorylated form. Phosphosites S12/13/T15, S27, S134/137/140 and S176/181 are highlighted in gold, and sites T55/56 and S116 in teal and labelled.

We next analysed Modanovo peptide predictions beyond the spectra identified by MaxQuant. For this, we set cutoffs on Modanovo confidence scores by leveraging the MaxQuant-identified PSMs as ground truth to map score cutoffs to precision estimates (Methods). As an initial positive control, we observed that, despite having no prior information on the sample preparation or enrichment strategy, Modanovo predominantly predicted unmodified peptides and peptides containing only common modifications (N-terminal acetylation or methionine oxidation) in the full proteome samples and phosphopeptides in the phospho-enriched samples for high precision estimates (Supplementary Fig. S14).

As an orthogonal proxy for peptide plausibility, we assessed the agreement between the experimental spectrum and the Prosit-predicted spectrum (42), using the spectral angle as a similarity metric (0 for no similarity, 1 for highest similarity, Methods). Since the currently available Prosit model does not handle PTMs, this analysis had to be restricted to unmodified peptides (and those containing only methionine oxidation). We stratified spectra by agreement between Modanovo and MaxQuant annotations. When considering all Modanovo predictions without applying any precision estimate, spectral angles were highest for sequences matching MaxQuant and lowest for spectra not identified by MaxQuant, presumably representing lower-quality spectra (Fig. 4B). A similar trend was observed when restricting to predictions with precursor mass agreement, although differences in median spectral angles were less pronounced. At the 90% precision threshold, Modanovo predictions matching MaxQuant showed high spectral similarity (median SA: 0.81), closely comparable to Modanovo-only predictions for spectra without MaxQuant identifications (median SA: 0.80; Fig. 4B). At 95%, spectral angles increased slightly across all groups, reaching 0.84 for both MaxQuant-matching and Modanovo-only predictions. For spectra where Modanovo and MaxQuant disagreed, spectral similarity also improved with confidence, with median SA values of 0.80 and 0.90 at the 90% and 95% thresholds, respectively (Fig. 4B). These results support the validity of many high-confidence Modanovo identifications and demonstrate that the model’s confidence score is an effective indicator of prediction quality. This is even evident for spectra not annotated by MaxQuant or where the two methods disagree, cases likely to be more challenging for peptide identification.

As a further source of evidence of the correctness of Modanovo identifications, we next evaluated the proportion of Modanovo-predicted peptides that perfectly aligned to the human or the MPXV proteome, allowing isoleucine-leucine mismatches (Methods). For PSMs from the phospho-enriched samples, the proportion of perfect alignments remained high, reaching 0.94 at an estimated precision of 95%, and 0.80 at 90% precision (Fig. 4C, Methods). In comparison, perfect alignment rates for the full proteome samples were slightly lower, reaching 0.84 and 0.72 at the 95% and 90% precision thresholds, respectively. Moreover, in both full proteome and phospho-enriched samples, PSMs corresponding to unmodified peptides or those containing only common modifications (methionine oxidation or N-terminal acetylation) exhibited consistently high alignment rates across the confidence range (Fig. 4D). For example, in the top 10,000 PSMs by confidence, the perfect alignment rate exceeded 93% and 77% for unmodified peptides on the full proteome samples and phospho-enriched samples, respectively (Fig. 4D). Notably, PSMs predicted to contain a phosphorylation event also maintained high perfect alignment rates, particularly in the phospho-enriched samples, where they reached a proportion of 0.93 perfect alignments in the top 10,000 PSMs and outperformed other PTM types across nearly the entire confidence spectrum. These results particularly support the reliability of phosphorylation predictions in the phospho-enriched samples.

We next assessed the novelty and complementarity of Modanovo’s predictions relative to database search by comparing the set of perfectly aligning peptides identified by Modanovo and MaxQuant to the reference proteomes with computed mass matching the experimental precursor, stratified by sample type and species of origin (human vs. MPXV; Fig. 4E). Notably, Modanovo identified a substantial number of MPXV peptides in their unmodified version that were missed by MaxQuant, particularly in the phospho-enriched samples. Specifically, Modanovo recovered 72 unique MPXV peptides in the phospho-enriched dataset that were not identified by MaxQuant, more than twice the number of peptides uniquely identified by MaxQuant (n=32), and 1,037 MPXV peptides in the full proteome dataset, compared to 239 unique to MaxQuant. While the number of shared peptides was higher for human than for viral sequences, this is expected given the higher abundance of host proteins and potential database coverage biases.

Allowing for alignment mismatches is particularly relevant in the context of viral proteomes, which often exhibit high sequence variability due to rapid mutation rates, strain differences, or incomplete annotation of viral coding sequences. When allowing for amino acid mismatches during alignment in the phospho-enriched samples, Modanovo’s advantage became more pronounced: the number of unique MPXV peptides increased by 129 and 247 when allowing for one and two mismatches, respectively (Supplementary Fig. S15). Moreover, aligning predicted peptides against a six-frame translation of the viral genome rather than the reference proteome alone revealed an additional set of unique MPXV peptides: 153 and 314 were identified by Modanovo under one-mismatch and two-mismatch conditions, respectively (Supplementary Fig. S15). These results highlight Modanovo’s strength in discovering peptides from viral proteins, particularly under phospho-enriched conditions where modified viral peptides may evade database search detection.

Moreover, we examined the organismal origin of Modanovo’s predictions by assessing the proportion of PSMs with perfect alignments to the human or MPXV proteome, stratified by PTM type and infection condition (Fig. 4F). Across all modification types, the majority of predicted peptides aligned to the expected species: human sequences appeared in both sample types (mock and MPXV-infected), while MPXV-specific peptides were observed almost exclusively in infected samples. Furthermore, to investigate temporal dynamics of host and viral peptide detection, we quantified the number of Modanovo-predicted PSMs from MPXV-infected samples with perfect alignment to the human or MPXV proteome across different time points post-infection (Fig. 4G). MPXV-derived peptides increased markedly over time. In contrast, human peptide counts remained relatively stable across all time points, regardless of modification type. Together, these results highlight Modanovo’s ability to recover biologically meaningful peptide sequences and capture dynamic changes in viral expression and post-translational modification patterns across infection time points.

Multiple seminal poxvirus studies highlighted the importance of phosphorylation dynamics during infection (35,48–51). We inspected phosphorylation sites identified in the MPXV multifunctional protein Cop-H5 (52). H5 was highlighted in the study that originally introduced this dataset as the most heavily phosphorylated viral protein with multiple known regulatory phosphosites (35). As an illustrative example, Modanovo consistently detected phosphosites at residues S12, S13, and T15 (cluster 1); S27; S134, S137, and S140 (cluster 2); S176, and S181; in agreement with those previously reported using MaxQuant (35). Phosphorylation of sites in cluster 2 and S176 was previously shown to regulate double-strand DNA binding activity, suggesting a dynamic role of this protein during the life cycle of this double-strand DNA virus (35). MaxQuant uniquely reported additional sites at S46 and T47, whereas Modanovo uniquely identified sites at T55, T56, and S116. One peptide was detected for sites T55 and T56 in two spectra, which were so far not described. For the target phosphosites at S116, Modanovo identified six distinct peptide sequences across 105 spectra, without a corresponding peptide reported by MaxQuant, predominantly at the later hours post-infection (12h and 24h, Supplementary Fig. 16). These PSMs showed high confidence scores and MS2 spectral evidence (Supplementary Fig. 17). The analogous phosphosite to S116 in the vaccinia virus (S109) in Cop-H5 has been reported previously as part of F10/H1 viral kinase/phosphatase phospho-network, a pivotal viral mechanism driving the phosphorylation dynamics during poxvirus lifecycle (51,53). These three phosphosites fall within a predicted N-terminal unstructured region of the protein, and to highly charged amino acid stretches thereof as inferred from AlphaFold structural (44) and electrostatic models (Fig. 4H). These three sites, uniquely identified by Modanovo, could thereby hint at functional sites of a Cop-H5 disordered region, potentially regulating its activities or di/multimerization potential. Taken together, these observations provide a proof-of-concept illustration of how DNPS can detect biologically relevant phosphosites and demonstrate how the approach can complement standard database search pipelines.

## Discussion

In this work, we curated a comprehensive dataset comprising a large number of peptide-spectrum matches (PSMs) from both in vivo experiments and synthetic peptides, including both unmodified peptides and post-translationally modified peptides containing 19 distinct amino acid-PTM combinations. This dataset enables model development and benchmarking of de novo peptide sequencing (DNPS) algorithms supporting modified peptides. Leveraging this dataset, we developed Modanovo, built by extending Casanovo to support 19 PTM-amino acid combinations within a single, unified model, without sacrificing performance on unmodified peptides. Modanovo achieved strong performance across a broad spectrum of modifications on the development dataset and on independent datasets, validating it as a robust and practical extension suitable for downstream applications. The application to a monkeypox virus (MPXV) dataset demonstrated the complementarity of Modanovo to a state-of-the-art database search approach, revealing hundreds of well-supported peptides missed by database search and revealing new MPXV phosphosites.

Modanovo was trained on synthetic peptides for most PTM types. Synthetic peptide data are less noisy and miss fewer fragment peaks than in vivo data from in vivo experiments. Nevertheless, these differences did not pose significant challenges during model transfer, underscoring the robustness of the learned representations. Nonetheless, the accurate prediction of longer, heavily modified peptides remains a known challenge for peptide identification tools and, especially, for autoregressive architectures, a limitation that could be addressed by future architectural refinements in the context of DNPS to modified peptides.

In addition to token-expansion approaches, including our work, Pi-PrimeNovo, and the recent model InstaNovo-P (54), one avenue for enhancement lies in the integration of open modification search tools, which could further expand PTM coverage without requiring explicit enumeration of all PTM-amino acid pairs during training. In this context, leveraging embeddings from chemical foundation models could enable representations that go beyond the mere residue masses, potentially resolving current ambiguities between residues of identical mass, such as isoleucine versus leucine, or O-GalNAc versus O-GlcNAc. In addition, coupling Modanovo with tools such as Prosit or data-driven rescoring pipelines such as Oktoberfest (55) may improve site localization by adding additional information such as retention times, particularly in spectra lacking strong site-determining fragment ions. The development and evaluation of such future tools could readily leverage the development dataset we provide, along with its splits into training, validation, and test sets.

In this study, we have applied Modanovo to datasets for which the reference proteome is known a priori. Remarkably, this still showed added value over database search. The advantages of DNPS are expected to be even more pronounced in scenarios where the reference proteome is poorly annotated, undergoes adaptive mutations, i.e., through selective pressure, when proteome sequence information is unavailable. These scenarios arise in studies of how RNA modifications affect protein sequence, in rapidly mutating tumours and RNA viruses, and in phospho-metaproteomics, where diverse microbial proteomes and widespread phosphorylation pose major challenges for database-driven approaches.

## Acknowledgments

We would like to thank the members of the Gagneur lab and Wilhelm lab for their valuable feedback, especially Johannes Hingerl, Joel Lapin, and Ludwig Lautenbacher. We are also grateful to members of the Kuster and Wilhelm lab for their support with data transfer, and to the Casanovo team for sharing their train/validation/test data splits.

## Data availability

No original data were created for this study. The raw files and MaxQuant identification files of the 21-PTM and the Monkeypox virus datasets were obtained through PRIDE accessions PXD009449 PXD040811, and PXD040889. The spectra and ground truth peptides of the MassIVE-KB datasets (v1 and v2) were downloaded from https://massive.ucsd.edu/ProteoSAFe/static/massive-kb-libraries.jsp. Spectral data and sequence annotations for the MULTI-PTM dataset were downloaded from Zenodo (56).

## Author contribution

Conceptualization: JG, MW, DKA; Data curation: DKA; Formal analysis: DKA, YB, CN, WG, VB; Funding acquisition: JG, MW; Investigation: DKA, YB, CN; Methodology: JG, MW, DKA; Project administration: JG, MW; Resources: JG; Software: DKA, YB, CN, WG; Supervision: JG, MW; Validation: DKA, VB, AP; Visualization: DKA, VB; Roles/Writing - original draft: DKA, JG, MW, VB, CN; Writing - review & editing: all authors.

## Abbreviations

AUPCC: Area under the precision-coverage curve
DNPS: De novo peptide sequencing
FDR: False discovery rate
HCD: Higher-energy collision dissociation
HFF: Human foreskin fibroblast
MPXV: Monkeypox virus
PTM: Post-translational modification
PSM: Peptide-spectrum match
SA: Spectral angle
m/z: mass-to-charge

## Supplementary Information

**Table 1:**
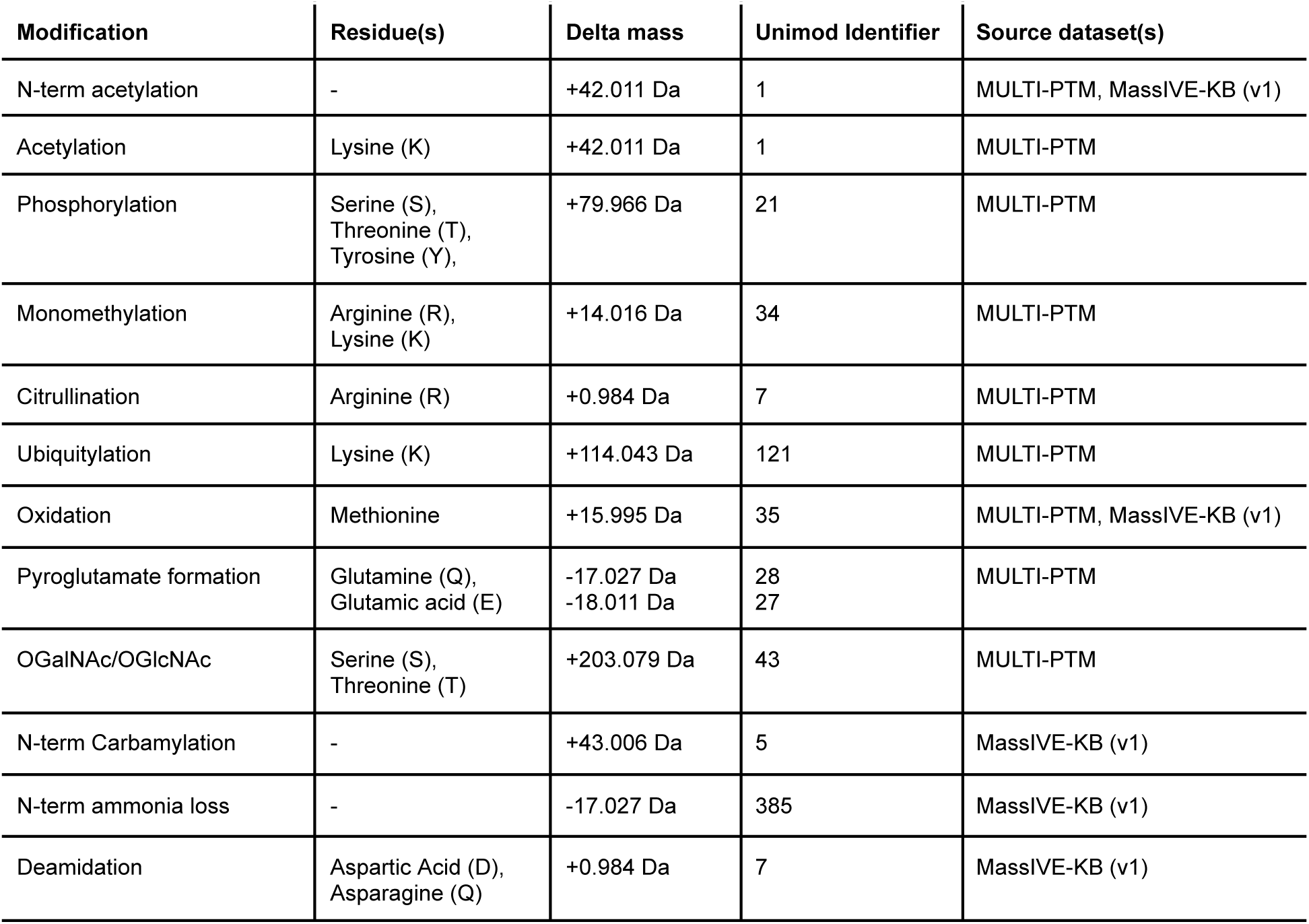
Post-translational modification (PTM)–amino acid combinations included in the development dataset. For each PTM, the amino acid that can contain this PTM, corresponding mass shifts (delta masses), and Unimod identifiers are provided, along with the source dataset(s) from which the modified peptides were obtained.

| Modification | Residue(s) | Delta mass | Unimod Identifier | Source dataset(s) |
| --- | --- | --- | --- | --- |
| N-term acetylation | - | +42.011 Da | 1 | MULTI-PTM, MassIVE-KB (v1) |
| Acetylation | Lysine (K) | +42.011 Da | 1 | MULTI-PTM |
| Phosphorylation | Serine (S),<br>Threonine (T),<br>Tyrosine (Y), | +79.966 Da | 21 | MULTI-PTM |
| Monomethylation | Arginine (R),<br>Lysine (K) | +14.016 Da | 34 | MULTI-PTM |
| Citrullination | Arginine (R) | +0.984 Da | 7 | MULTI-PTM |
| Ubiquitylation | Lysine (K) | +114.043 Da | 121 | MULTI-PTM |
| Oxidation | Methionine | +15.995 Da | 35 | MULTI-PTM, MassIVE-KB (v1) |
| Pyroglutamate formation | Glutamine (Q),<br>Glutamic acid (E) | -17.027 Da<br>-18.011 Da | 28<br>27 | MULTI-PTM |
| OGalNAc/OGlcNAc | Serine (S),<br>Threonine (T) | +203.079 Da | 43 | MULTI-PTM |
| N-term Carbamylation | - | +43.006 Da | 5 | MassIVE-KB (v1) |
| N-term ammonia loss | - | -17.027 Da | 385 | MassIVE-KB (v1) |
| Deamidation | Aspartic Acid (D),<br>Asparagine (Q) | +0.984 Da | 7 | MassIVE-KB (v1) |

**Supplementary Figure S1:**
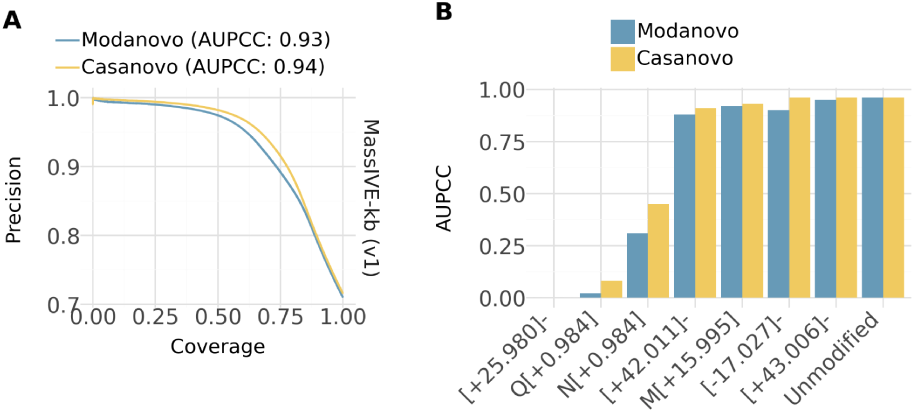
Performance on the MassIVE-KB (v1) dataset compared to Casanovo. A, Precision-coverage curves at the peptide level comparing Modanovol (blue) to Casanovo (v4, yellow) on the test set of the MassIVE-KB (v1) dataset, originally used for training Casanovo, consisting mostly of unmodified peptide sequences. **B**, Area under the precision-coverage curve (AUPCC) obtained using Modanovo (blue) and Casanovo (v4, yellow) for the different PTM-residue combinations contained in the test set of MassIVE-KB (v1).

**Supplementary Figure S2:**
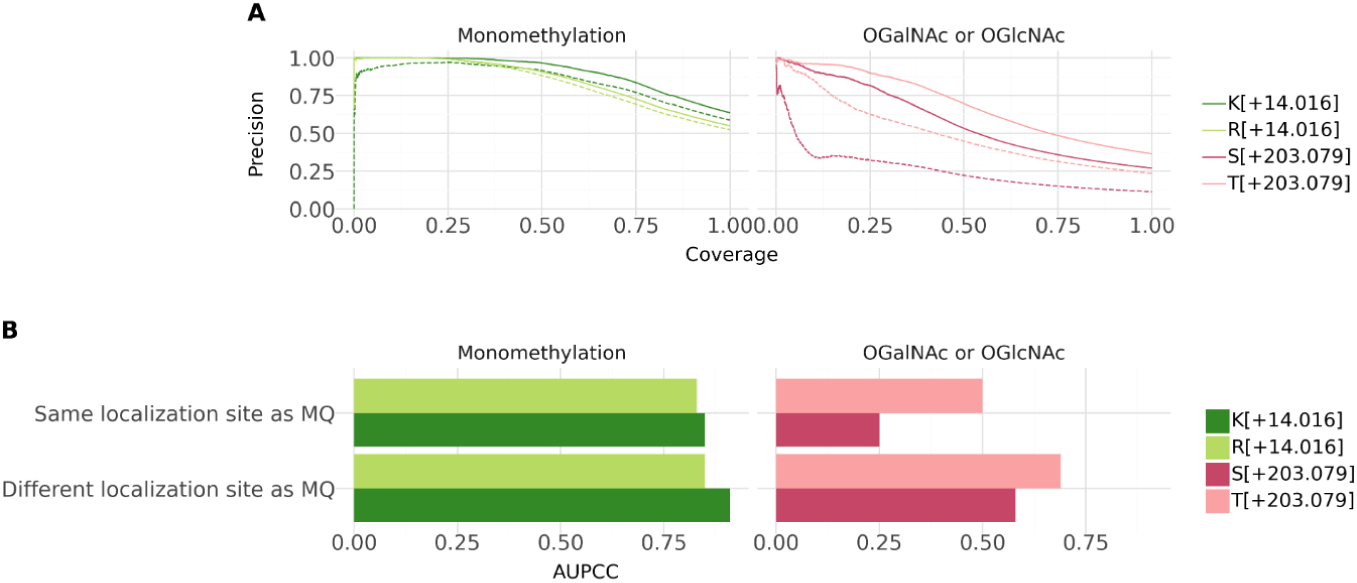
Performance on monomethylation, O-GalNAc, and O-GlcNAc modifications allowing for different modification sites. A, Precision-coverage curves at the peptide level for the PTM types monomethylation and OGalNAc/OGlcNAc in the MULTI-PTM dataset. Curves are shown separately for lysine (K[+14.016]), arginine (R[+14.016]), serine (S[+203.079]), and threonine (T[+203.079]). Solid lines indicate cases where the predicted sequences are considered correct, even in cases where the predicted PTM localization site differs from the site reported by MaxQuant (MQ), while dashed lines indicate peptide correctness only when the same localization sites as MaxQuant were predicted. **B,** Area under the precision-coverage curve (AUPCC) for lysine (K[+14.016]), arginine (R[+14.016]), serine (S[+203.079]), and threonine (T[+203.079]) for cases where the predicted sequences are considered as correct, even in cases where the predicted PTM localization site differs from the site reported by MaxQuant (MQ), and for peptide correctness only when the same localization sites as MaxQuant were predicted.

**Supplementary Figure S3:**
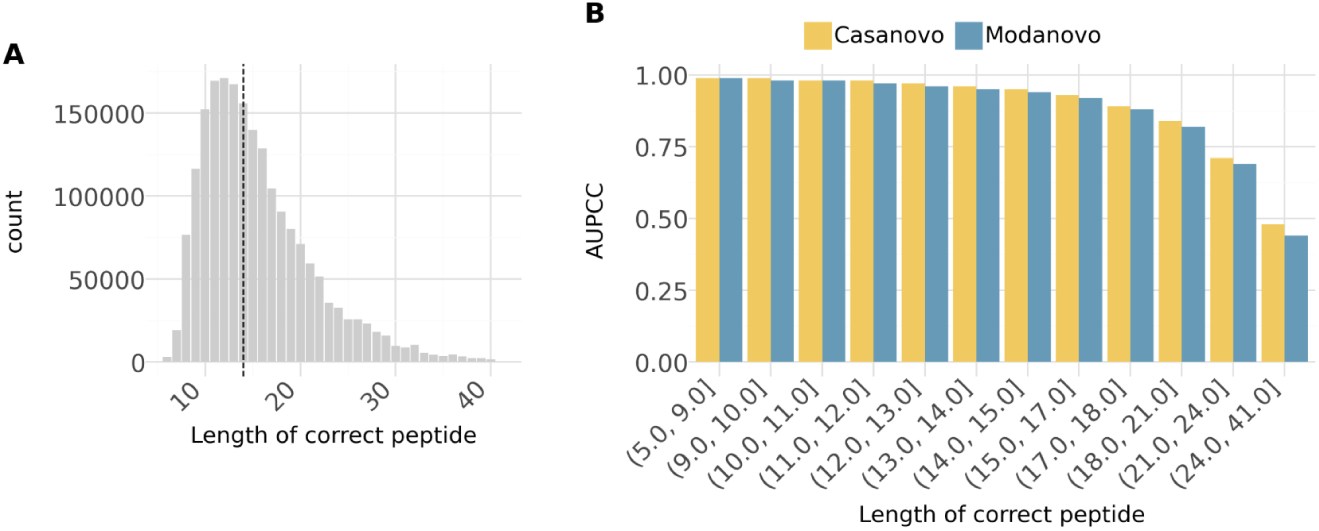
Performance by peptide length on the MassIVE-KB (v1) dataset. **A**, Distribution of peptide lengths in the MassIVE-KB (v1) dataset. Dashed vertical lines indicate median peptide length. **B,** Area under the precision-coverage curve (AUPCC) for different peptide length categories obtained with Casanovo (yellow) and Modanovo (blue).

**Supplementary Figure S4:**
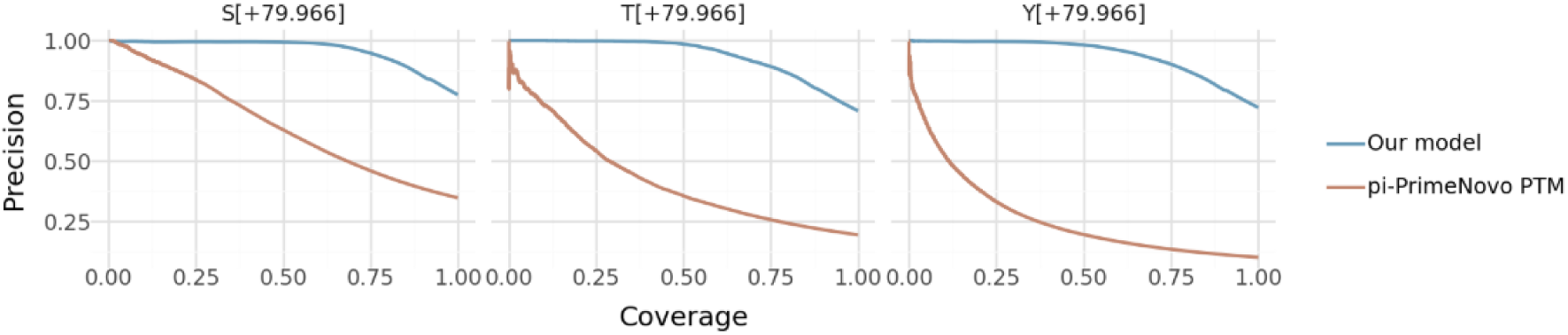
Performance on phosphorylated peptides in comparison to π-PrimeNovo-PTM. Precision-coverage curves at the peptide level for phosphorylation in the MULTI-PTM dataset. Curves are shown separately for serine (S[+79.966]), threonine (T[+79.966]), and tyrosine (Y[79.966]) for Modanovo (blue) and the π-PrimeNovo-PTM model (brown), which was fine-tuned to allow the prediction of phosphorylated residues.

**Supplementary Figure S5:**
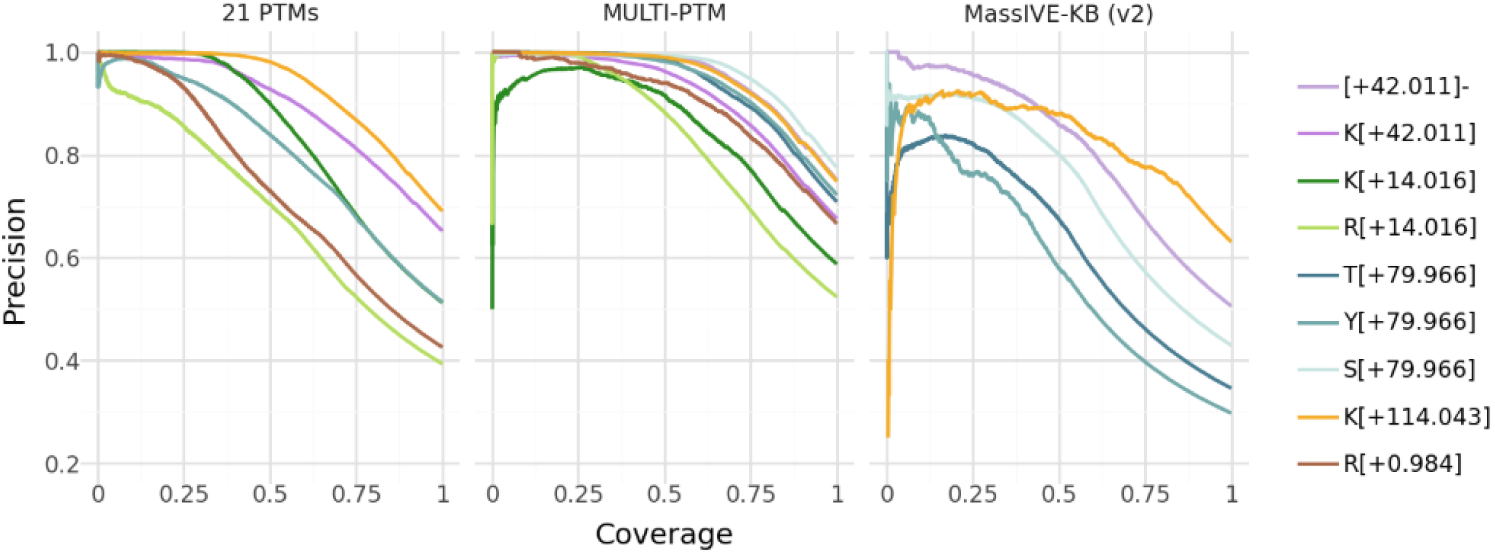
Precision-coverage curves for Modanovo’s predictions at the peptide level for the different PTM-residue combinations in the three different datasets, MassIVE-KB (v2), MULTI-PTM and 21 PTMs from ProteomeTools. PTM-residue combinations are restricted to those seen during model training and overlapping with those in MassIVE-KB (v2) and the 21-PTM dataset.

**Supplementary Figure S6:**
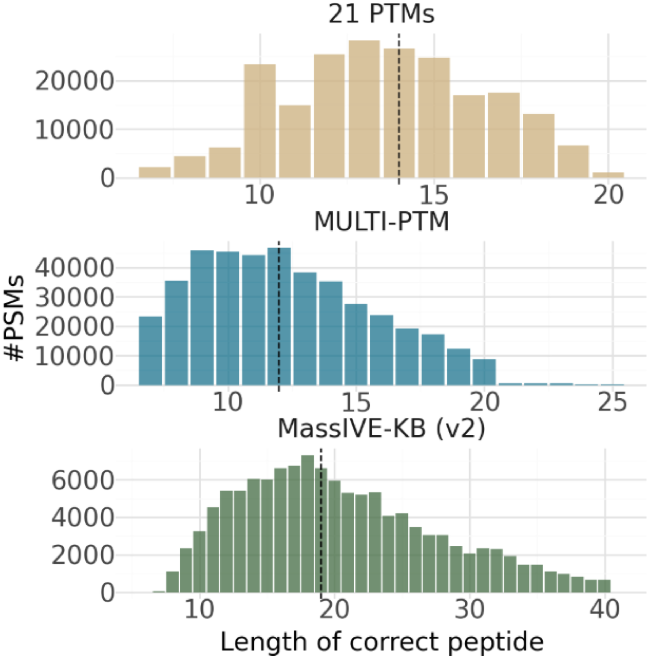
Peptide length distribution in the datasets MassIVE-KB (v2) and 21 PTMs. The top panel shows the number of peptide-spectrum matches (PSMs) as a function of peptide length for the 21-PTM dataset (bottom panel), the MULTI-PTM dataset (middle panel), and the MassIVE-KB (v2) dataset (bottom panel). Dashed vertical lines indicate the median peptide length in each dataset. PSMs restricted to those carrying modifications seen during model training and overlapping with those in MassIVE-KB (v2) and the 21-PTM dataset.

**Supplementary Figure S7:**
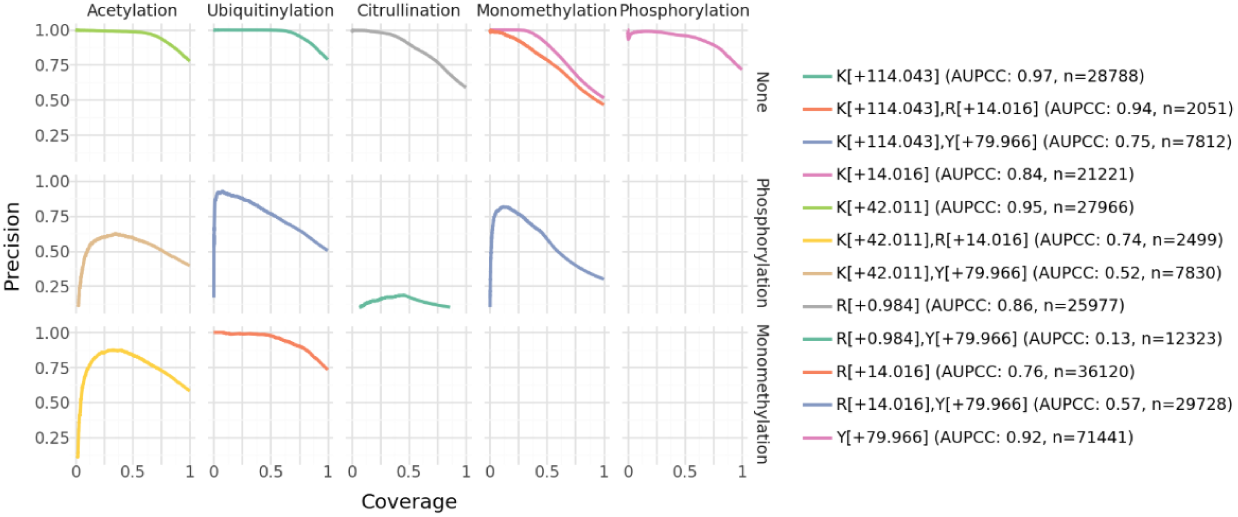
Performance on the 21-PTM dataset, including peptides with multiple PTM types. Precision-coverage curves are shown for peptide sequences containing either a single PTM type (first row) or combinations of two PTM types (second and third rows; e.g., acetylation and phosphorylation in the second row, first column). Colored lines indicate specific amino acid-PTM combinations, with the corresponding area under the precision-coverage curve (AUPCC) reported in the legend.

**Supplementary Figure S8:**
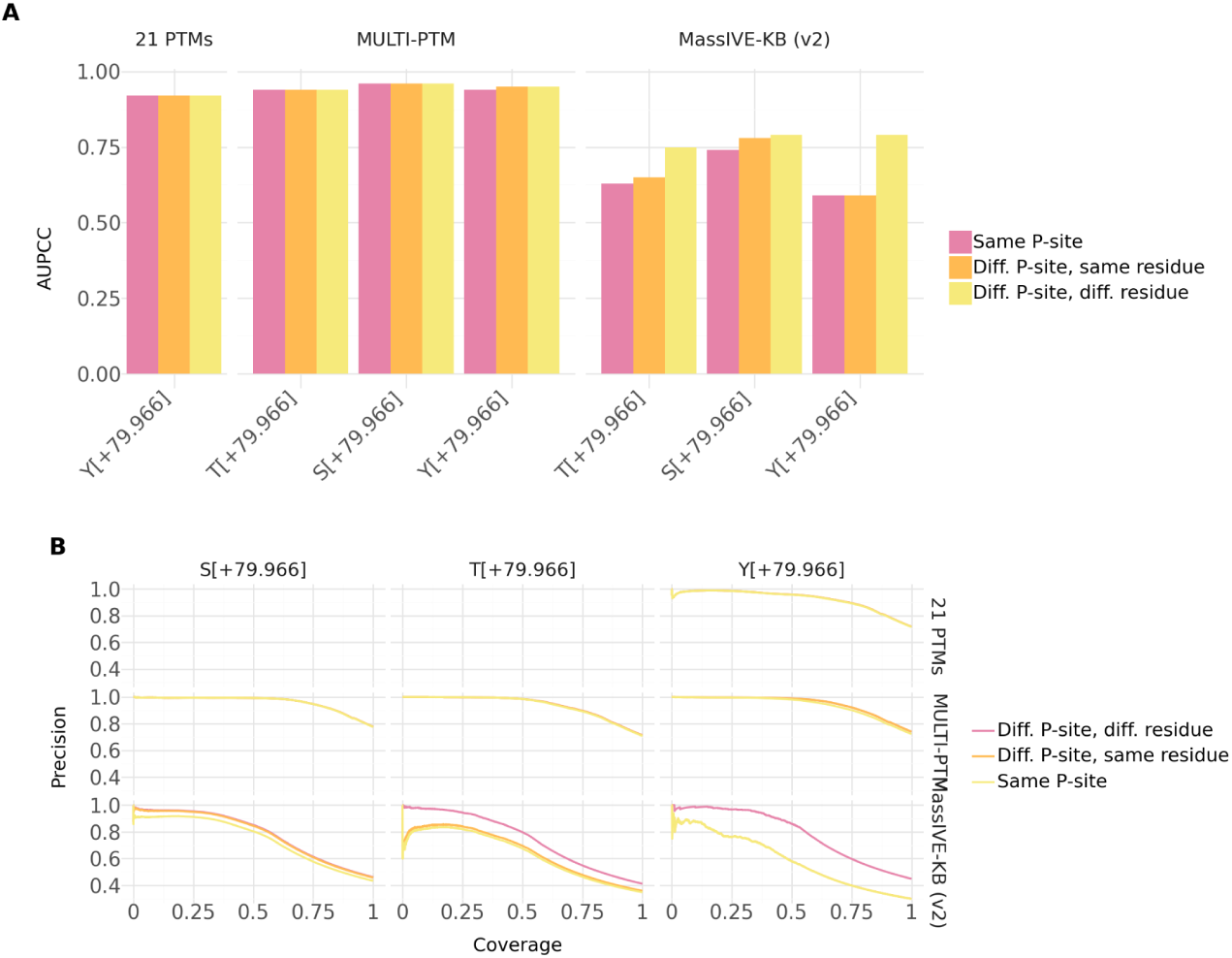
A, Area under the precision-coverage curve (AUPCC) for the different phosphorylated residues on the MassIVE-KB (v2), MULTI-PTM and 21-PTM datasets, comparing against the same P-site as the ground truth peptide (rosa), allowing for a different P-site between Modanovo’s predictions and the ground truth peptide on the same residue (orange) and allowing for a different P-site on a different residue (yellow). **B**, Same as A, but displaying the precision-coverage curves for the different residues and P-site considerations.

**Supplementary Figure S9:**
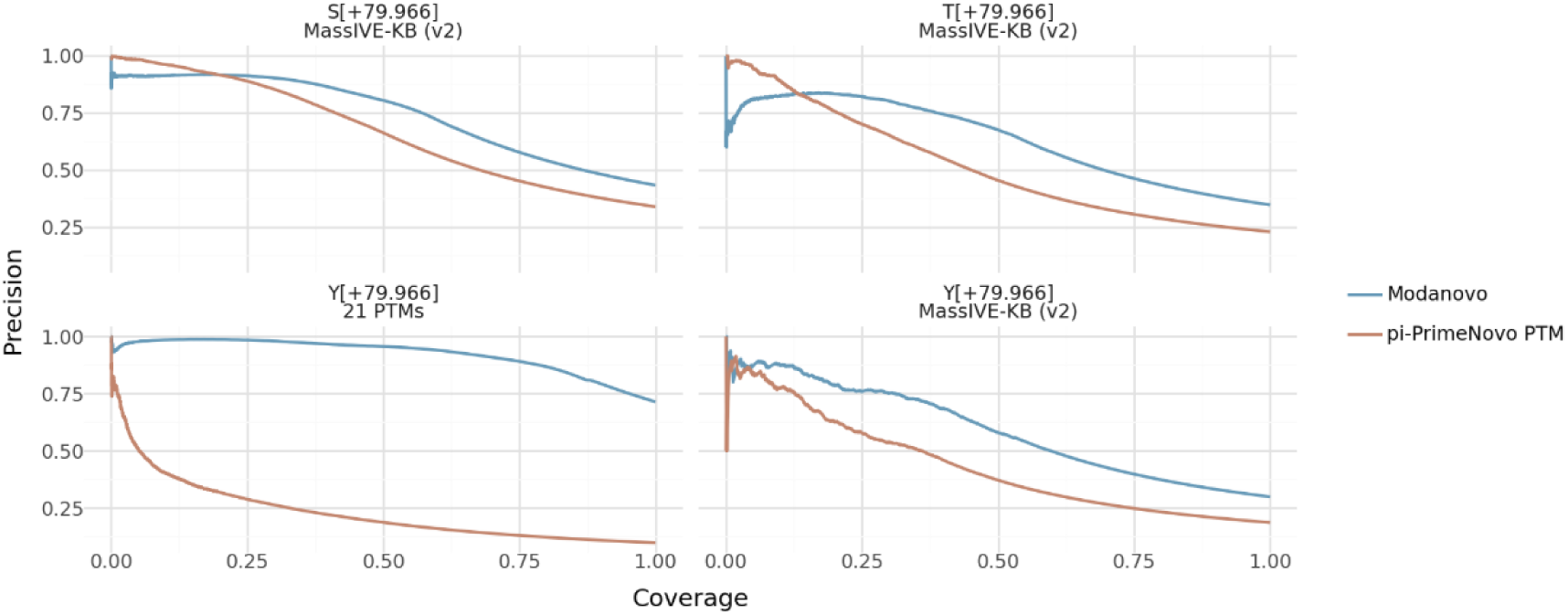
Phosphorylation performance on the MassIVE-KB (v2) and 21-PTM datasets compared to π-PrimeNovo-PTM. Precision-coverage curves for phosphorylation on serine (S), threonine (T), and tyrosine (Y) residues (+79.966 Da) for Modanovo (blue) compared to π-PrimeNovo-PTM (brown) fine-tuned to predict phosphorylated residues on the 21-PTMs and MassIVE-KB (v2) datasets (facets).

**Supplementary Figure S10:**
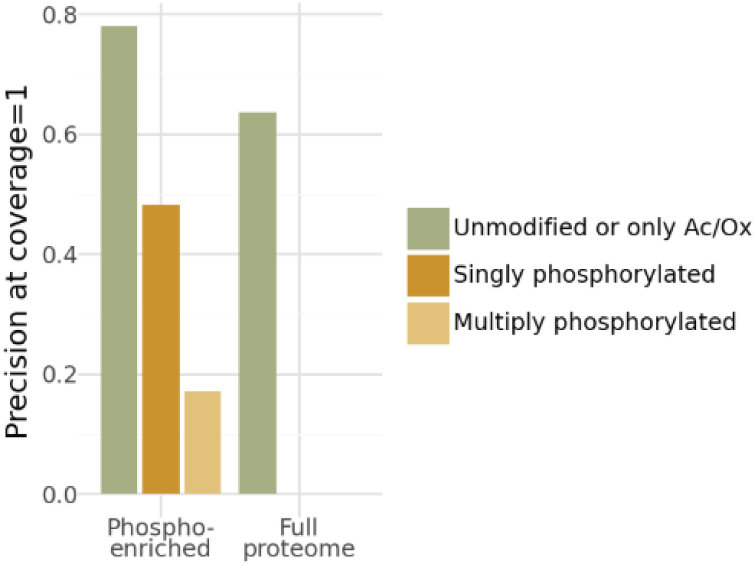
Performance on the MPXV dataset compared to MaxQuant identifications. Final precision (at coverage=1) obtained by Modanovo with MaxQuant peptides as ground truth sequences for comparison for the full proteome samples and phosphorylation-enriched samples, with the MaxQuant sequence having multiple phosphorylated residues (light mustard), one phosphorylation residue (mustard), and no phosphorylation residues (green).

**Supplementary Figure S11:**
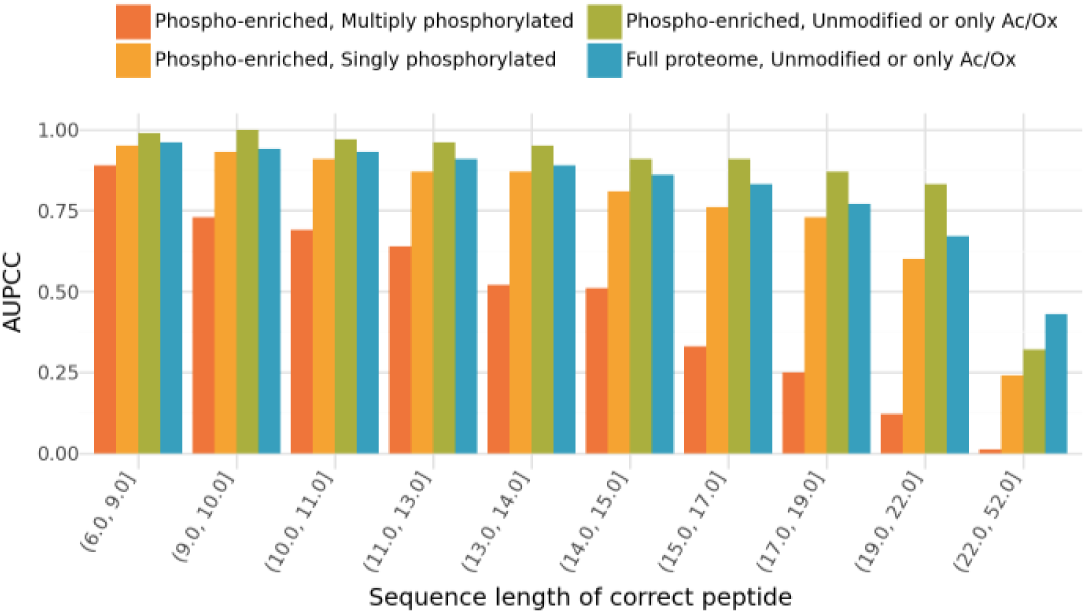
Modanovo’s performance on the MPXV dataset stratified by different peptide length categories. Area under the precision-coverage curve (AUPCC) by Modanovo on the MPXV dataset, stratified by peptide length category and modification status: multiply phosphorylated peptides (orange), singly phosphorylated peptides (yellow), unmodified peptides or those containing only N-terminal acetylation/oxidation in the phospho-enriched dataset (green), and unmodified peptides or those containing only N-terminal acetylation/oxidation in the full proteome dataset (blue).

**Supplementary Figure S12:**
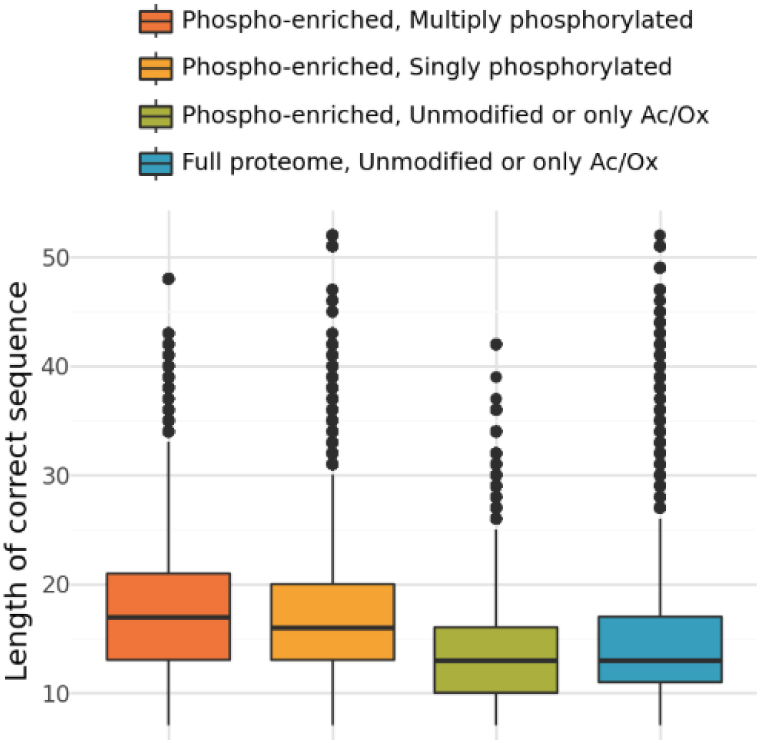
Peptide length distributions of MaxQuant peptides on the MPXV dataset. Distribution of peptide lengths of the MaxQuant peptides in the MPXV dataset for multiply phosphorylated peptides (orange), singly phosphorylated peptides (yellow), unmodified peptides or those containing only N-terminal acetylation/oxidation in the phospho-enriched dataset (green), and unmodified peptides or those containing only N-terminal acetylation/oxidation in the full proteome dataset (blue).

**Supplementary Figure S13:**
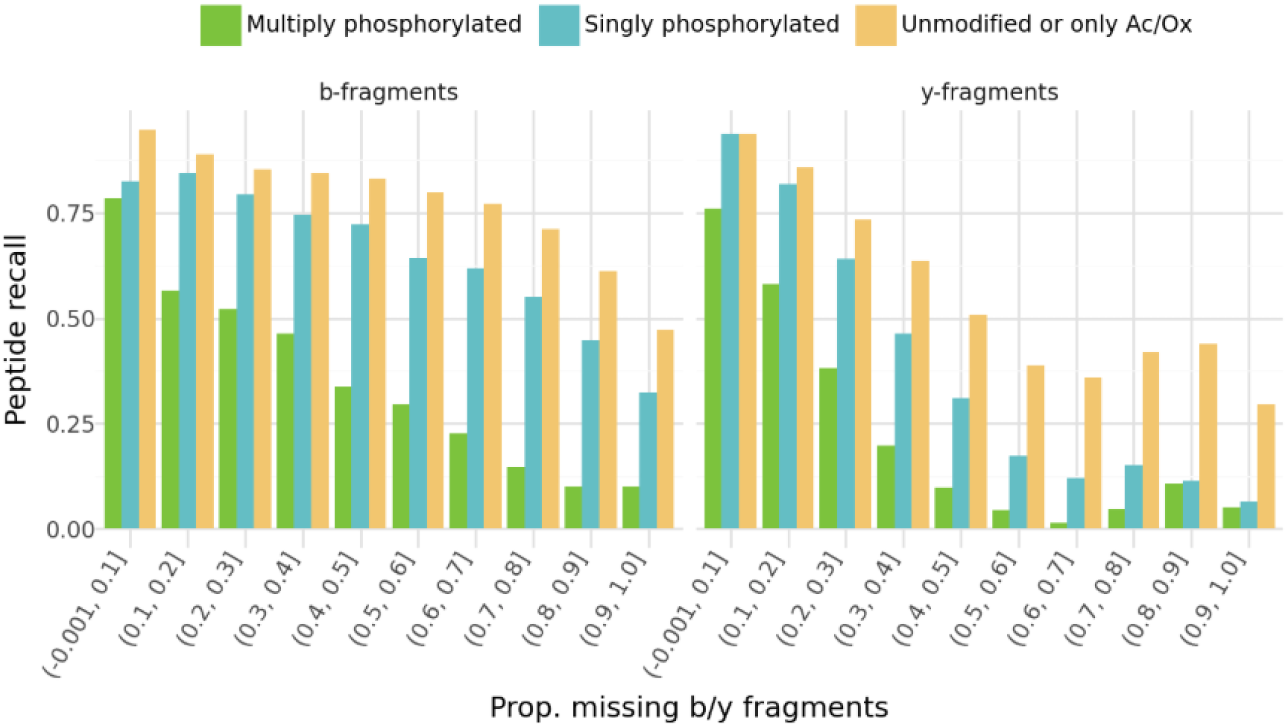
Impact of fragment ion coverage on model performance in the MPXV dataset. Final precision (at coverage=1, i.e., peptide recall) as a function of the proportion of missing b-ions (left) and y-ions (right) in the spectra, stratified by peptide type: multiply phosphorylated (green), singly phosphorylated (blue), and unmodified or containing only N-terminal acetylation/oxidation (yellow) in the phospho-enriched samples of the MPXV dataset.

**Supplementary Figure S14:**
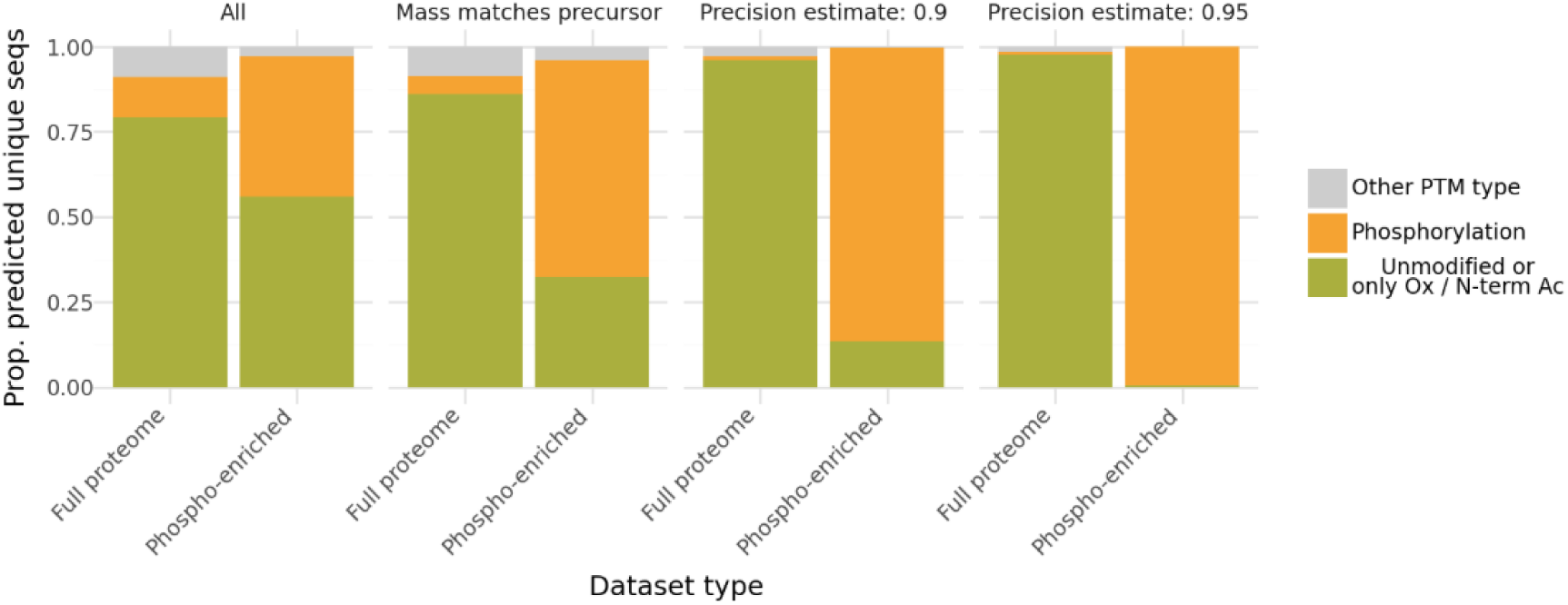
Modanovo’s predictions on the MPXV dataset. Proportion of predicted unique peptide sequences in the MPXV dataset by dataset type (full proteome vs. phospho-enriched) and different filtering criteria: all predictions, mass-matched to precursor, and subsets with estimated precision ≥0.9 and ≥0.95. Bars show the relative contribution of unmodified peptides or those with only oxidation/N-terminal acetylation (green), phosphorylated peptides (orange), and peptides with other PTM types (gray).

**Supplementary Figure S15:**
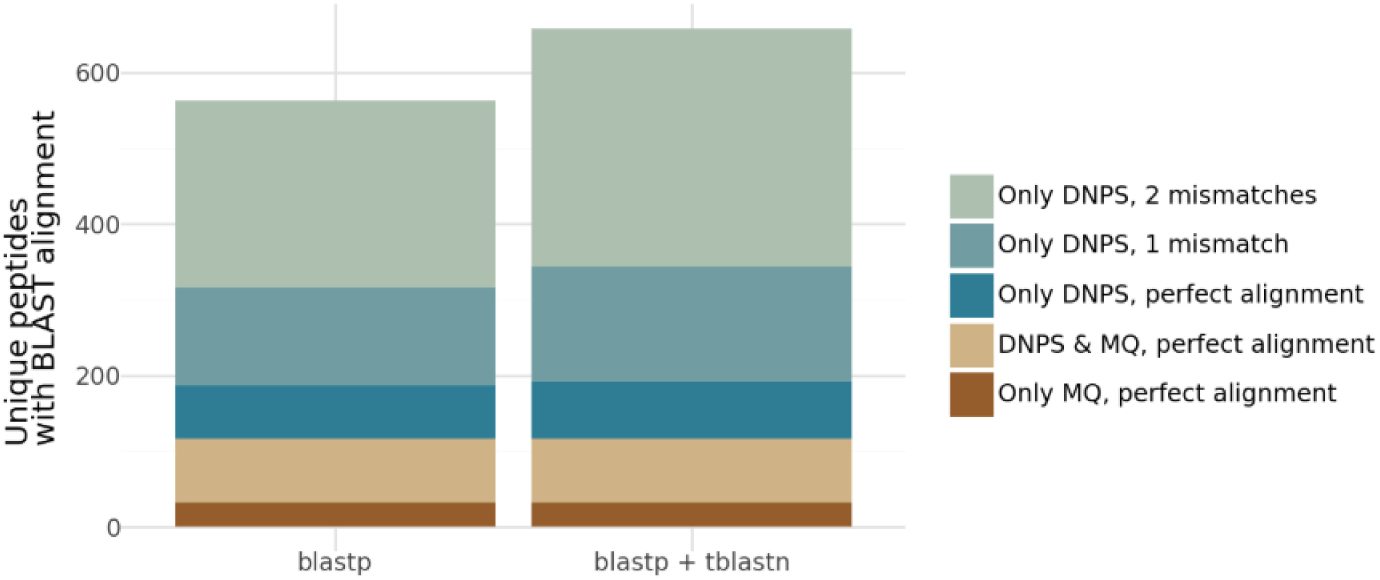
Comparison of unique peptide identifications in the MPXV dataset based on BLAST alignment. Number of unique peptides recovered using blastp alone (left) or blastp combined with tblastn (right), stratified by source of identification: only DNPS with two mismatches (light green), only DNPS with one mismatch (teal-green), only DNPS with perfect alignment (blue), peptides identified by both DNPS and MaxQuant (MQ) with perfect alignment (beige), and only MQ with perfect alignment (brown).

**Supplementary Figure S16:**
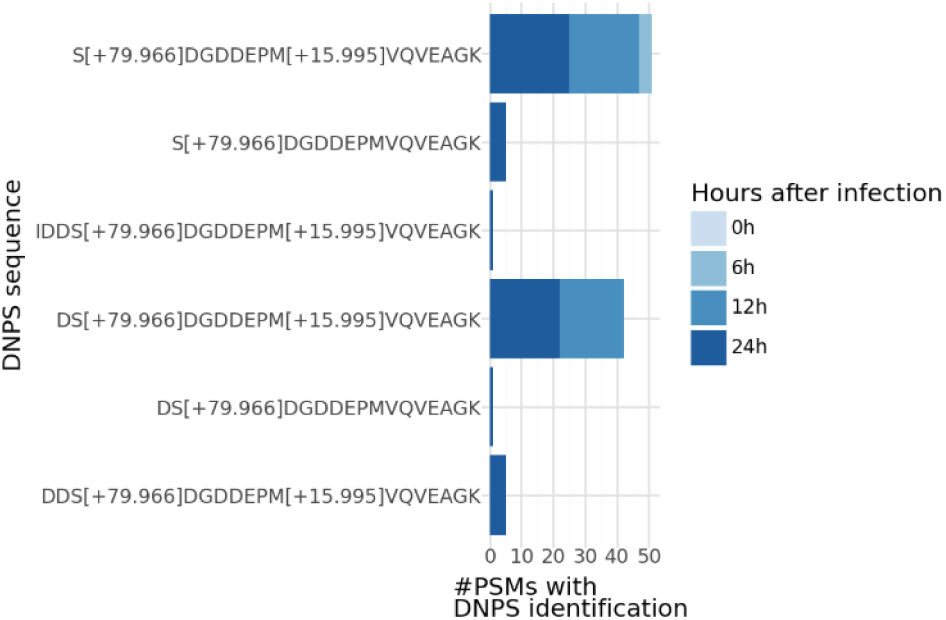
Temporal detection of DNPS-identified peptides carrying the H5 phosphosite S116. Total number of PSMs identified for each peptide sequence containing the S116 phosphorylation site in the viral protein H5, separated by infection time point (0, 6, 12, and 24h).

**Supplementary Figure S17:**
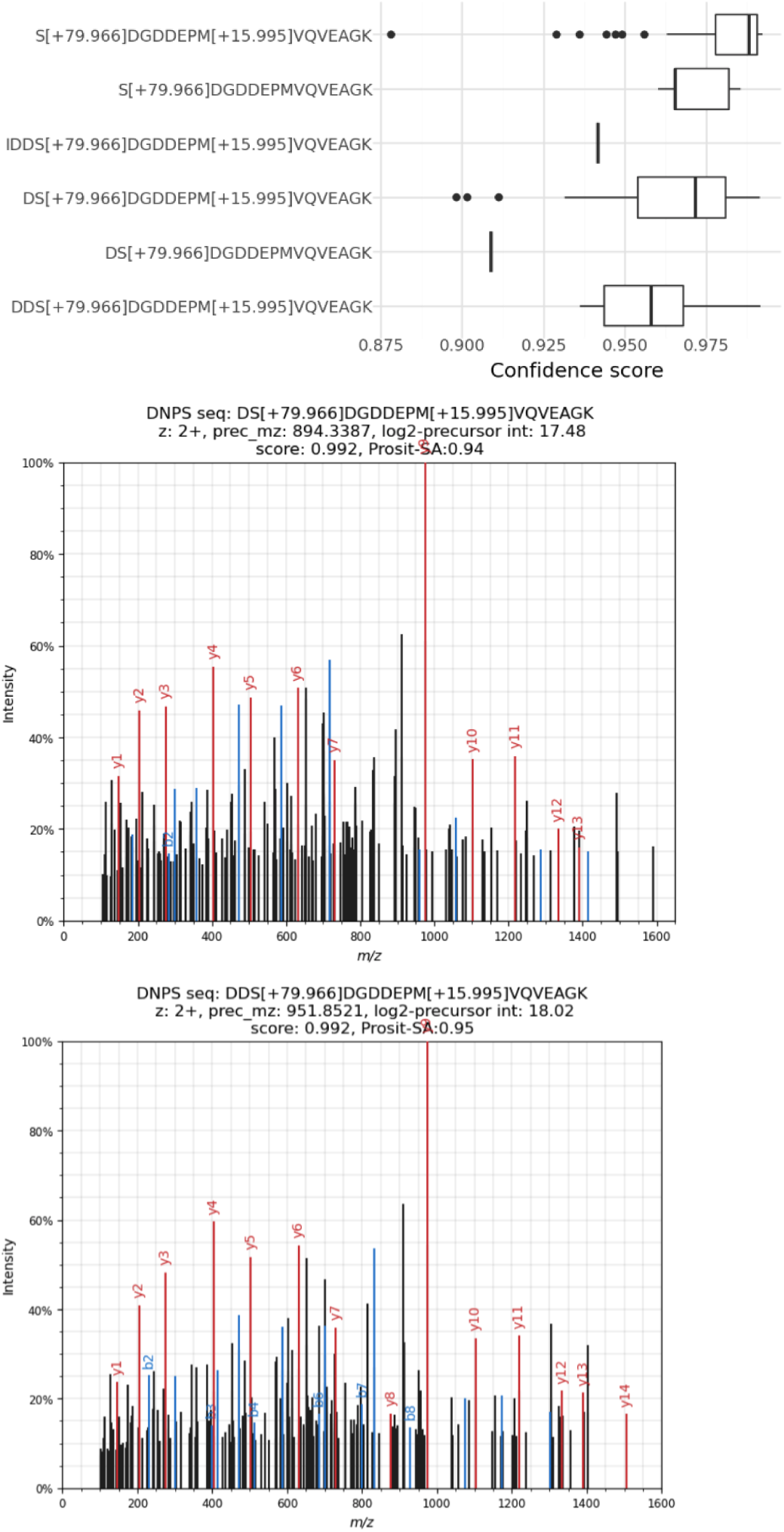
MS2 spectral evidence for the Modanovo-identified phosphosite S116 in H5. A, Modanovo’s confidence scores for the six distinct peptide sequences carrying the S116 phosphorylation site in the viral protein H5, each supported by multiple PSMs. **B,** Example annotated spectrum for the peptide DS[+79.966]DGDDEPM[+15.995]VQVEAGK, illustrating fragment ion coverage around the phosphorylated serine. **C,** Same as B, but for the peptide sequence DDS[+79.966]DGDDEPM[+15.995]VQVEAGK.

